# Meet me in the middle: brain-behavior mediation analysis for fMRI experiments

**DOI:** 10.1101/2020.10.17.343798

**Authors:** Jules Brochard, Jean Daunizeau

## Abstract

Functional outcomes (e.g., subjective percepts, emotions, memory retrievals, decisions, etc…) are partly determined by external stimuli and/or cues. But they may also be strongly influenced by (trial-by-trial) uncontrolled variations in brain responses to incoming information. In turn, this variability provides information regarding how stimuli and/or cues are processed by the brain to shape behavioral responses. This can be exploited by brain-behavior mediation analysis to make specific claims regarding the contribution of brain regions to functionally-relevant input-output transformations. In this work, we address four challenges of this type of approach, when applied in the context of mass-univariate fMRI data analysis: (i) we quantify the specificity and sensitivity profiles of different variants of mediation statistical tests, (ii) we evaluate their robustness to hemodynamic and other confounds, (iii) we identify the sorts of brain mediators that one can expect to detect, and (iv) we disclose possible interpretational issues and address them using complementary information-theoretic approaches. *En passant*, we propose a computationally efficient algorithmic implementation of the approach that is amenable to whole-brain exploratory analysis. We also demonstrate the strengths and weaknesses of brain-behavior mediation analysis in the context of an fMRI study of decision under risk. Finally, we discuss the limitations and possible extensions of the approach.

## Introduction

Functional outcomes (e.g., subjective percepts, emotions, memory retrievals, decisions, etc…) are partly determined by external stimuli and/or contextual cues. But they may also be strongly influenced by irreducible variability in brain responses to incoming information (Ferster, 1996; Shadlen and Newsome, 1994). In particular, neural noise may be a critical determinant of illusory percepts, aberrant emotions, erroneous memory retrievals, biased decisions, etc… (Bays, 2014; Faisal et al., 2008; Hong and Rebec, 2012). For most existing statistical data analyses of neurophysiological data, neural noise is typically treated as a statistical nuisance, since it compromises the identification of relationships between measured brain activity and experimental variables (Doi and Lewicki, 2011; Naselaris et al., 2011). This perspective is unfortunate however, since neural noise can provide complementary information regarding how incoming information is processed and/or distorted by the brain to yield functional outcomes (Dinstein et al., 2015; McDonnell and Ward, 2011; Stein et al., 2005). The critical point is that a brain system may encode functionally-relevant information that is not *used* by the brain when producing a functional outcome. This has been repeatedly demonstrated in neurological patients who do not exhibit significant behavioral impairments despite being lesioned in brain regions that are known to encode behaviorally-relevant information(Aerts et al., 2016; Alstott et al., 2009). But what if one can show that neural noise contributes to -otherwise unexplained-behavioral variability? This is the essence of *brain-behavior mediation analysis*, which aims at detecting neural systems that both respond to behaviorally-relevant cues or stimuli and eventually impact overt behavior (MacKinnon et al., 2007).

Recall that any cognitive function can be seen as some form of -potentially complex, context-dependent, redundant, partially unconscious, etc-neural transformation of relevant stimuli into adaptive behavioural outcomes (Robbins, 2011). By adaptive, we simply mean that cognitive functions serve a specific purpose, which can be abstracted and put to a (behavioural) test. At the limit, one could argue that understanding cognitive functions reduces to assessing input-output relationships, where inputs are experimentally controlled stimuli and/or task instructions, and outputs are overt behavioural outcomes. In this view, neuroimaging in healthy subjects should serve to identify how brain networks contribute to the input-output transformation (Palestro et al., 2018; Rigoux and Daunizeau, 2015; Turner et al., 2019). A reasonable strategy here is to identify intermediary neural states that *mediate* the impact of incoming information onto overt behavior and/or subjective reports. In its simplest form, brain-behavior mediation analysis reduces to a twofold regression analysis that aims at detecting uncontrolled variability in brain responses that significantly improves behavioral predictability. The ensuing statistical tests typically reason as follows: if region M responds to experimental factor X, and explains behaviour Y above and beyond the effect of X, then M mediates the effect of X onto Y. For example, brain-behavior mediation analysis was used to identify the prefrontal and/or subcortical systems that mediate successful emotional regulation (Wager et al., 2008), threat response (Wager et al., 2009a, 2009b) or risk avoidance (Yamamoto et al., 2015). More recently, the anterior cingulate cortex, the anterior insula, the thalamus and some brain stem nuclei were shown to mediate various aspects of pain perception (Atlas et al., 2010, 2014; Geuter et al., 2018; Koban et al., 2017, 2019; Woo et al., 2015). Most of these studies were performed using the multilevel mediation/moderation or M3 toolbox (Wager, 2008), which was first derived for probing effective connectivity from fMRI signals. Since then, a few multivariate extensions of brain-behavior mediation analysis were proposed, aiming at improving either spatial or temporal resolution (Chén et al., 2018; Lindquist, 2012; Zhao and Luo, 2017). But these approaches neither lay out nor address the specific methodological and interpretational challenges posed by brain-behavior analysis, when applied to typical fMRI experiments. In our view, progress in brain-behavior mediation analysis requires answering at least four important (and related) questions:

- (Q1) Which test statistics should be used? Not only should the test statistics be valid (i.e. yield controlled false positive rate), but they also should be maximally powerful. The latter is a pressing issue because fMRI induces a massive multiple comparison problem, which can only be solved by using more stringent significance thresholds (Lindquist and Mejia, 2015; Worsley and Friston, 1995). We will summarize and compare the statistical properties of the most established test statistics of mediation analysis.
- (Q2) How robust is brain-behavior mediation analysis to assumptions regarding the hemodynamic response function (HRF) and other confounds? Recall that virtually all forms of fMRI time-series analyses rely on HRF models to assess effects of interest (Deshpande et al., 2010; Gitelman et al., 2003; Liao et al., 2002; Pedregosa et al., 2015). Although brain-behavior mediation analysis involves similar assumptions, different modelling strategies may be employed that yield distinct bias-variance tradeoffs. We will compare the statistical properties of these candidate approaches in the presence of deviations to modelling assumptions.
- (Q3) What sort of brain mediators can we expect to detect? Consider the bottom-up chain of neural information processing stages that eventually yield behavioral outcomes (from low-level sensory processing to high-level cognitive treatment of stimuli and/or cues). It turns out that these stages do not have the same chance of being detected. As we will see, this is a corollary consequence of the nontrivial (and yet undisclosed) impact of neural noise onto the statistical properties of mediation analysis.
- (Q4) Does mediation analysis induce potential interpretational issues? As we will see, some interpretational issues are specific to the chosen statistical testing approach, but others are generic to any brain-behavior mediation analysis. In particular, significant mediated effects are compatible with two distinct scenarios regarding the causal relationship between brain activity and behavioral responses. We discuss the importance of this and related issues and identify ways to address them.

In this work, we address these four questions from a user-oriented statistical perspective. Our aim here is to set a methodological standard for brain-behavior mediation analysis. The Methods section serves as the statistical and conceptual basis for addressing the four questions (Q1-Q4) above. It starts with a description of the brain-behavior mediation model and its associated null-hypothesis testing alternatives. Specific issues that arise in the context of typical fMRI experiments (factorial designs and condition contrasts, group-level random effects analysis, etc) are shortly discussed. We then consider the critical role of neural noise in brain-behavior mediation analyses, and present alternative solutions to the issue of HRF deconvolution. We close this section with a note on causality and its accompanying interpretational issue. We address the latter using a complementary information-theoretic approach (so-called *I/O test*). *En passant*, we show how to exploit the underlying mathematical degeneracy to drastically reduce the computational cost of whole-brain mediation analysis. In the Results section, we use numerical Monte-Carlo simulations to answer questions Q1-Q4. We compare the specificity and sensitivity of candidate mediation tests, as a function of neural noise, and in the presence of hemodynamic confounds. We also evaluate the utility and robustness of our I/O test. We then strengthen our *in-sillico* conclusions with an application to an experimental fMRI dataset acquired when people make decisions under risk. We exemplify the use of brain-behavior mediation analysis to ask questions regarding intra- and between-subjects variations in behavioral responses and attitudes. Finally, we discuss our results in the light of the existing literature and highlight potential weaknesses and perspectives (Discussion section).

## Methods

In what follows, we will consider behavioral paradigms akin to decision tasks, whereby subjects need to process some (experimentally-controlled) information *X* to provide a (measured) behavioral response *Y*. Brain-behavior mediation analysis then aims at identifying whether some (anatomically-specific) feature of their observed brain activity *M* mediates the effect of *X* onto *Y*. In our example fMRI application (see Results section), we will focus on a value-based decision making task, whereby participants have to accept or reject (response *Y*) a risky gamble composed of a 50% chance of winning a gain G and a 50% chance of loosing L (input information *X*). But more generally, *X* is an experimental manipulation of some sort, *M* is a measure of neural activity at the time of processing the stimulus, and *Y* is some overt expression of the stimulus-induced covert mental state of interest.

### The brain-behavior mediation model

Let *n* be the number of trials in a typical experimental session. Let *X*, *M* and *Y* be *n* × 1 column vectors encoding the trial-by-trial experimental manipulation, the brain’s response to the experimental manipulation (e.g., the magnitude of the fMRI BOLD response to the stimulus at each trial, in some voxel or region of interest) and the behavioral response to the experimental manipulation, respectively. For the sake of mathematical simplicity, and without loss of generality, we will assume that *X*, *M* and *Y* have all been z-scored.

From the perspective of identifying the determinants of behavior, one may first ask whether *X* has an effect on *Y* or not. In its simplest mathematical form, this question reduces to considering the following simple linear regression model:

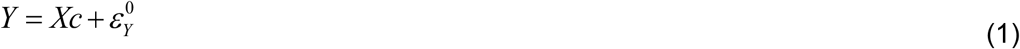

where *c* is an unknown regression coefficient that measures the strength of the statistical relationship between the independent (*X*) and dependent (*Y*) variables, and 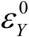 are model residuals. On would then simply test for the statistical significance of *c*, under some assumptions regarding the distribution of model residuals 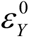.

Now, one may also ask whether *M* mediates the effect of *X* onto *Y*. In its simplest mathematical form, this question relies on the following pair of linear regression models:

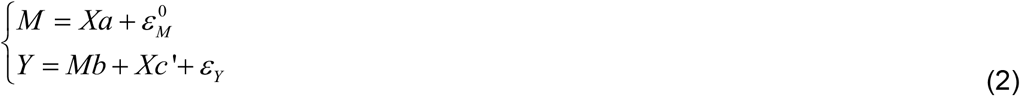

where the first equation expresses the fact that *M* responds to *X* (with some unknown susceptibility *c′*), and the second equation states that *Y* depends upon both *M* (with some unknown susceptibility *b*) and *X* (with some unknown susceptibility *c*′). On may think of residuals 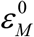 in terms of some form of *neural noise*, because they capture trial-by-trial variations in *M* that are independent of *X*. As we will see, they play a pivotal role in brain-behavior mediation analysis.

Although simple, Equation 2 does not explicitly quantify the size of a mediated effect. But this can be done by noting that Equation 2 can be rewritten as follows:

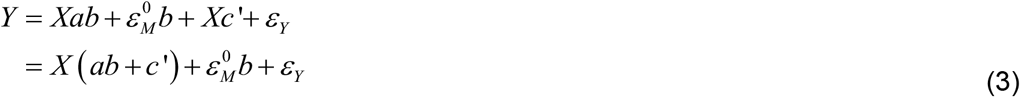

where *M* has simply been replaced by its expression from Equation 2. Equation 3 is helpful in realizing that the *total* effect of *X* onto *Y* is partitioned into a *direct* effect (whose size is *c*′) and an *indirect* effect (whose size is *ab*). This distinction is important because the latter is the effect of *X* onto *Y* that is mediated by *M*. This is why established mediation tests rely on assessing the statistical significance of the indirect effect (MacKinnon et al., 2007). Note that so-called *full mediation* occurs when *c′* =0 (no direct path), and one speaks of *partial mediation* whenever *c′* = 0.

Importantly, when we perform mass-univariate mediation analysis, we effectively consider each voxel or region of interest in isolation, and ask whether the local indirect effect is statistically significant. If mediation tests are repeated over voxels, then they form a statistical mediation *map*, which can localize which brain structure(s) mediate(s) the effect of *X* onto *Y*. In this context, Equations 1–3 have two interesting implications, which we will highlight now.

To begin with, recall that the incoming information *X* is processed by a distributed brain system, whose elements (sampled across large voxel sets) concurrently contribute to the behavioural response *Y*. The structure of this distributed brain system is likely to involve multiple processing pathways that work both in series and in parallel, as in Figure 1 below.

**Figure 1:**
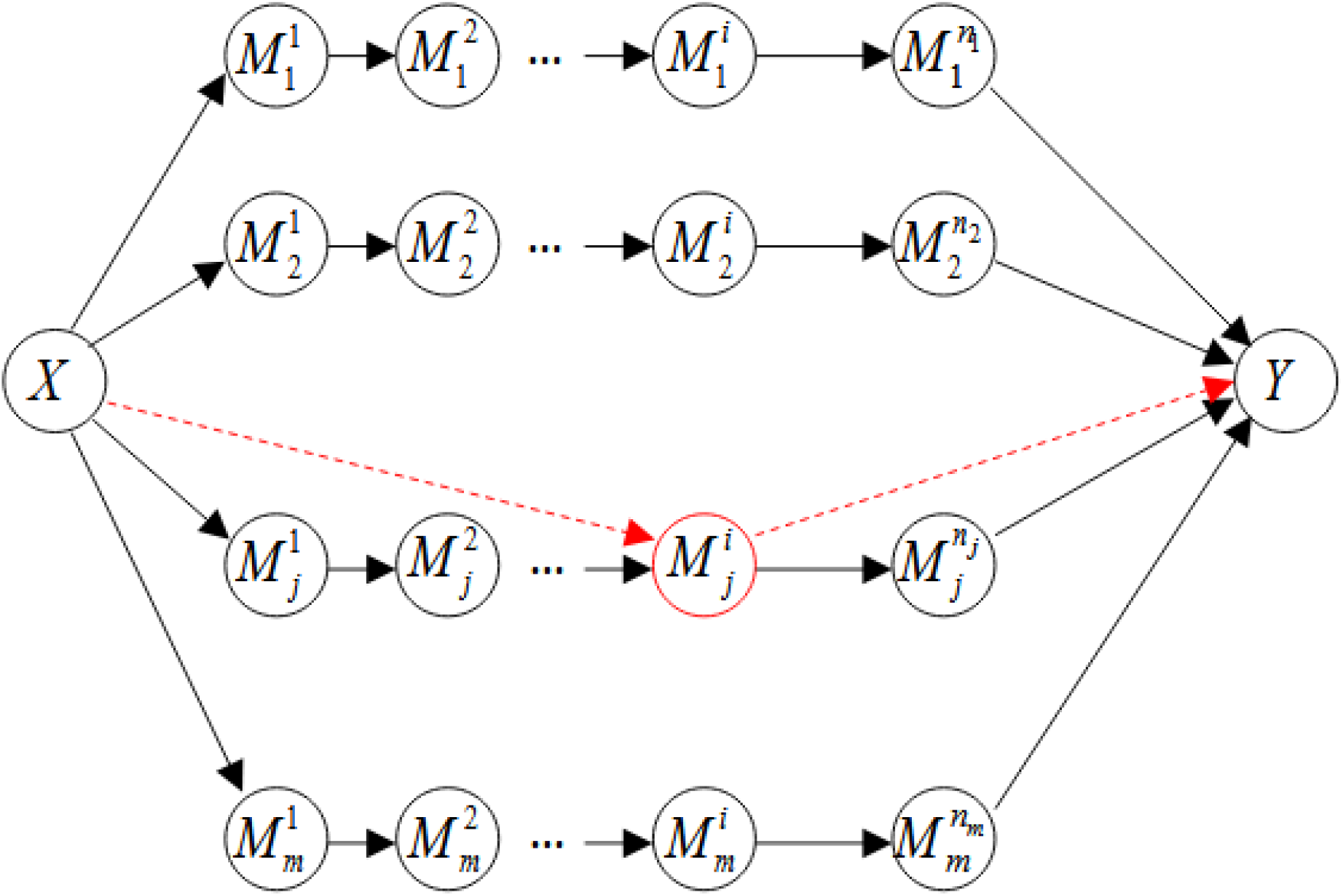
Example structure of a processing hierarchy in the brain.M. Here, *X* and *Y* encode the experimental manipulation and the ensuing behavioral response, respectively. Variables 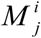 are activity within brain regions that act as intermediary processing steps. In this oriented graphical model, arrows represent causal relationships. Although processing pathways operate both in series and in parallel, mass-univariate brain-behavior mediation analysis ignore this and treat each region/voxel independently of each other (red dotted arrows).

In simple bottom-up hierarchical architectures such as this one, lower levels would correspond to e.g., occipital low-level visual processes, whereas higher levels would map to, e.g., prefrontal decision making processes. Clearly, Figure 1 is already an oversimplification because it ignores reciprocal connections, branching processes and/or context-dependent gating mechanisms (Friston, 2011; He and Evans, 2010; Rubinov and Sporns, 2010). But when we perform mass-univariate brain-behavior mediation analysis, we reduce the complexity even further by considering each voxel or region of interest in isolation, effectively ignoring any hierarchical structure of this sort.

First, given the likely parallel nature of processing pathways, one would not expect that any isolated voxel or region of interest may ever *fully* mediate the impact of *X* onto *Y*. The implicit assumption of mass-univariate brain-behavior mediation analysis is that, in each voxel, the direct path *c*′ effectively captures, in a non specific manner, mediated effects that go through other (parallel) pathways. This, however, places a very heavy load on the statistical sensitivity of mediation tests, which need to be able to detect potentially small indirect effect sizes, even when correcting for multiple comparisons (e.g., across voxels).

Second, nothing prevents different processing pathways to have strong but opposing impacts on the behavioral response. An example here would be opponent brain systems that yield strong but ambivalent (e.g., appetitive-aversive) cognitive states, whose idiosyncratic balance may explain one’s specific ability to suppress e.g., impulsive behavioral responses (Seymour et al., 2005; Zhang et al., 2017). In particular, if the impact of different pathways balance out, then the total effect of *X* onto *Y* may become undetectable (*c ≈* 0). It follows that brain-behavior mediation analyses may be required for faithfully identifying the determinants of behavior. Alternatively, the relative contribution of different pathways may vary across individuals, which may drive inter-individual behavioral differences. We will see an example of this in the Results section below.

### Statistical tests of mediation

In what follows, we recall the most established approaches to null-hypothesis testing of mediated effects. We start with the premise that if *M* mediates the effect of *X* onto *Y*, then the corresponding indirect effect has to be different from 0 (*ab* ≠0). In what follows, we summarize two kinds of statistical testing approaches (namely: the “indirect” and the “conjunctive” approaches) that differ in terms of how they frame the corresponding null hypothesis.

The *indirect* approach follows from noting that the null hypothesis of mediation analysis can be framed as follows: 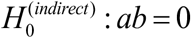.

Under the simple brain-behavior mediation model in Equations 1–2, the indirect effect equates the difference between total and direct effects, i.e. *ab* = *c* − *c′*. This is why early approaches to mediation testing were assessing the statistical significance of the difference *c−c*’ (Baron and Kenny, 1986). However, theoretical work demonstrated that this equivalence may not always hold (Pearl, 2012), which would render the ensuing test invalid. This applies to typical fMRI experiments, because of the effect of confounding variables on path coefficient estimates.

Another, more valid, approach is to compare estimates of the indirect effect to their distribution under the null. This is the principle of *Sobel’s test* (Sobel, 1982). Recall that all parameters are identifiable from Equation 2, given *X*, *Y* and *M*. In particular, the ordinary least-squares (OLS) estimates 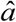 and 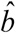 of unknown path coefficients *a* and *b* are given by (all *X*, *Y* and *M* variables are z-scored, see Appendix A for a mathematical derivation):

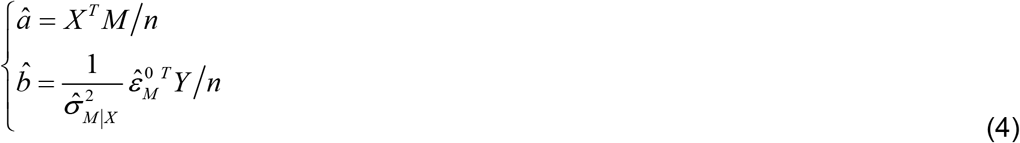

where the neural noise estimate 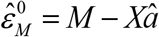 is the component of *M* that cannot be explained by *X*, and 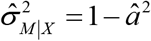 is its sample variance. From Equation 4, one can see that 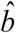 is simply the sample correlation between behavioral data *Y* and the neural noise estimate 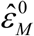. In other words, 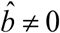 when *M* has an effect on *Y above and beyond the effect of X*. In addition, the variance of these OLS estimates are given by:

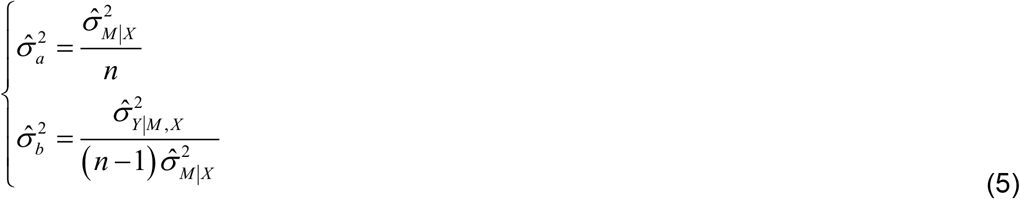

where 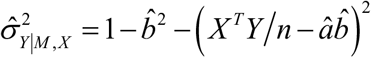 is the sample variance of behavioral residuals’ estimates 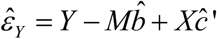.

Under the assumption that model residuals *ε_Y_* and 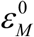 are i.i.d. normal variables, then both 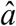 and 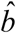 follow normal distributions: 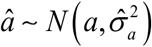 and 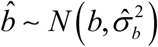 It can then be shown (see Appendix B) that the product 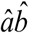 approximately follows a normal distribution, i.e.: 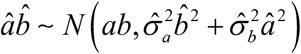. This implies that, under the null, the following pseudo z-statistics:

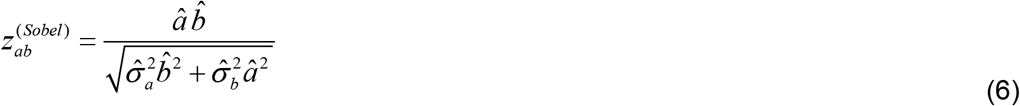

approximately follows a Student probability density function. This then serves to derive p-value of Sobel’s unsigned (two-tailed) significance test 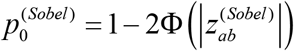, where φ is Student’s cumulative density function with appropriate degrees of freedom.

Later improvements over Sobel’s test (Hayes and Scharkow, 2013) derived from theoretical statistical works on the distribution of the product of two normal random variables, which essentially include an additional 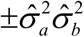 term to the denominator of Sobel’s pseudo-zscore (Aroian, 1947; Goodman, 1960).

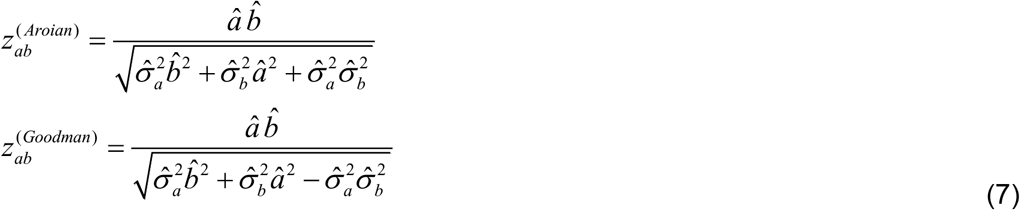

We refer to these extensions as Aroian’s and Goodman’s tests, respectively. Alternatively, non-parametric approaches have been proposed to derive the distribution of indirect effect size estimates under the null (MacKinnon et al., 2002, 2004). Here, we will use the same bias-corrected bootstrap approach as the one proposed in the M3 toolbox (Wager, 2008).

The *conjunctive* approach follows from noticing that the null hypothesis of mediation analysis is composite (Moran, 1970), i.e.: 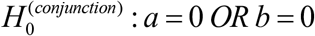. Of course, both null hypotheses are exactly equivalent, but the composite null highlights the fact there is no mediated effect as long as one path coefficient is null (which breaks the causal cascade). In turn, one may test for the conjunction of both effects, i.e. test for the statistical significance of both *a* and *b* path coefficients. In practice, *conjunctive* testing relies on the “maximum p-value” approach (here, two-tailed test):

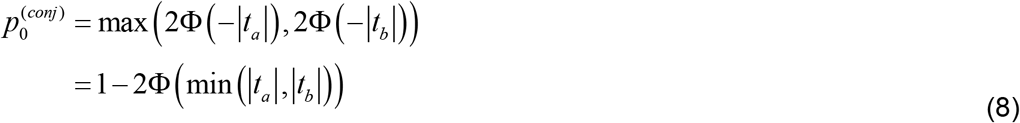

where 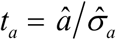 and 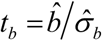 are Student’s test statistics of *a* and *b* path coefficients, respectively. Formally speaking, 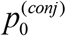 provides an *upper bound* on the joint probability that, under the null, two independent Student’s test statistics take more extreme values than *t_a_* and *t_b_* (Friston et al., 2005a; Nichols et al., 2005). This is important, because conjunctive testing cannot be invalid but may have low sensitivity. However, it is trivial to show that Sobel’s pseudo z-score is always smaller than the conjunctive test statistics, i.e.: 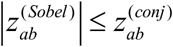, where 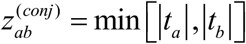 is the conjunctive test statistics (see Appendix B). This means that one would expect conjunctive testing to be systematically more efficient than Sobel’s approach. At this point, we note that the sensitivity profile of indirect and conjunctive approaches actually depends upon neural noise strength and model misspecifications (see next sections). We will address this and related issues in the Results section, using extensive numerical Monte-Carlo simulations.

We refer the reader interested in extending these statistical approaches to experimental designs including multiple conditions (cf., e.g., factorial designs) and/or multiples subjects (cf. group-level random effects analysis) to Appendix C and D, respectively.

### The non-trivial impact of neural noise

Although indirect and conjunctive null hypotheses are formally equivalent to each other, the latter is helpful to disclose the subtle tension behind mediation testing. In brief, two conditions must be satisfied for detecting a mediated effect: (i) strong evidence for *a* ≠ 0 and (ii) strong evidence for *b* ≠ 0. The former means that *X* partly explains the trial-by-trial variability of *M*. And the latter means that *M* partly explains the variability of *Y* that is unexplained by *X*. The critical point here is to realize that these two conditions are in conflict with each other. This is because they have opposing demands on neural noise 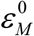. Note that the conjunctive test statistics 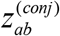 is given by:

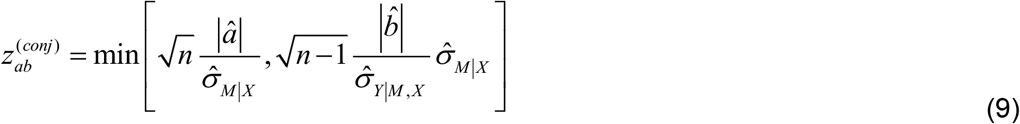

where we simply have inserted Equation 5 into the definition of the conjunctive test statistics. One can see that the standard deviation 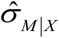 of the neural noise estimate 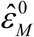 will have opposing effects on the conjunctive test statistics. In brief, if 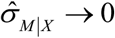, then 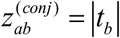, which tends towards 0 when 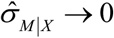. Recall that, by definition, 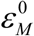 is the component of *M* that cannot be explained by *X* (cf. Equation 2). Thus, in the absence of neural noise, the evidence for *a* ≠ 0 is maximal, but *M* cannot explain any variability in *Y* that is unexplained by *X*, i.e. the evidence for *b* ≠0 is minimal. Reciprocally, if 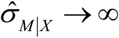, then 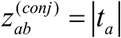, which tends also towards 0 when 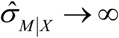. In other words, if neural noise strength is very high, then evidence for *a* ≠ 0 is weak. Only for *intermediary levels of neural noise* can evidence for both *a* ≠ 0 and *b* ≠ 0 reach statistical significance. We note that this observation generalizes to any mediation test, irrespective of the mathematical form of the brain-mediation model. We refer the interested reader to Appendix E.

We will quantify the impact of neural noise on the statistical efficiency of candidate mediation testing approaches in the Results section below. But this property of mediation analysis has an important implication, which we now highlight.

Recall the structure of the processing hierarchy in Figure 2. Within a given processing pathway, each hierarchical level responds to its (lower-level) parents, eventually changing the information content in an incremental manner, e.g.:

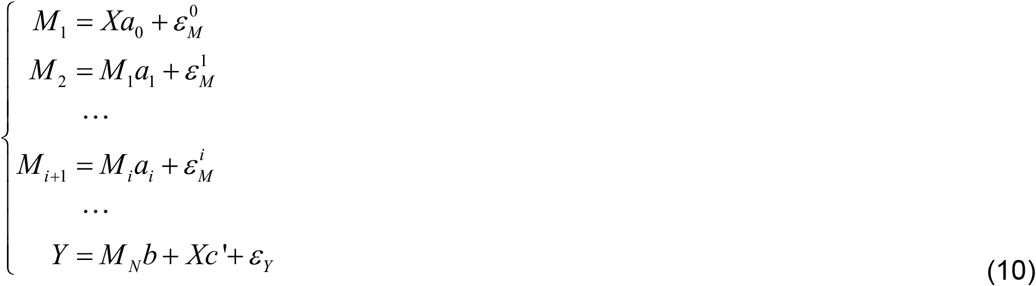

where {*M*_1_,*M*_2_,…,*M_i_*,*M_N_*} are local neural responses (indexed by their level along the hierarchy), and local neural noise increments 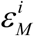 effectively capture, in an agnostic manner, the unique contribution of each hierarchical level. Here, one would expect that local neural responses gradually diverge from the initial explanatory variable *X*. This is simply because the correlation between *X* and the local neural response *M_i_* degrades as the accumulated neural noise increments 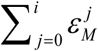 increases. In turn, one would expect that mass-univariate mediation analysis can only detect those neural information processing steps that are positioned at an intermediary hierarchical level, i.e. sufficiently far away from either end of the hierarchy. We will exemplify this in the Results section below.

**Figure 2:**
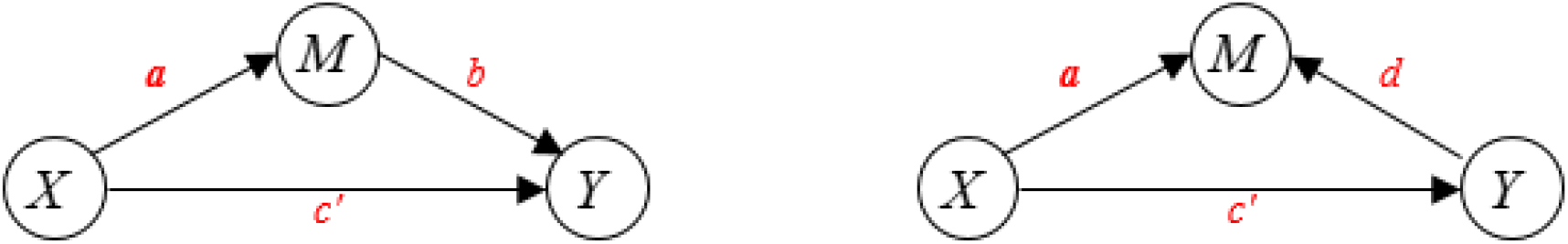
The two causal interpretations of mediated effects. Left panel: “native” causal interpretation of brain-behavior mediation analysis (cf. Equation 2). Right panel: “swapped” causal interpretation of brain-behavior mediation analysis (cf. Equation 11). Corresponding path coefficients are shown in red.

### Dealing with hemodynamic confounds

Clearly, the brain-behavior mediation model in Equation 2 cannot directly be applied to fMRI time series. The reason is twofold. First, behavioral and neural variables are not sampled in the same manner. In brief, the former is collected at each “trial” of the behavioral task, while the latter is typically sampled at a sub-trial temporal resolution. Second, fMRI BOLD dynamics effectively result from the convolution of neural activity with the hemodynamic response function or HRF (Logothetis et al., 2001; Martin et al., 2006). This implies that the event-related BOLD response is delayed in time, when compared to trial onsets. In addition, if the inter-trial interval is smaller than the HRF duration (which is typically the case), BOLD signals measured during a trial may derive from the additive contributions of multiple neural responses (to the current and preceding trials). For the purpose of brain-behavior mediation analysis, there are essentially two ways of dealing with such hemodynamic confounds.

On the one hand, one may deconvolve BOLD signals from the HRF, as follows. Let *τ_k_* (resp., Δ_*k*_) be the onset time (resp., duration) of the *k* th trial in the experimental design. One first construct “trial” regressors that span the duration of the fMRI session (at the sampling resolution of fMRI ; typically: TR=1-2secs), which are zero everywhere except during the time interval defined as [*τ_k_*, *τ_k_* + Δ_*k*_]. Each of these is then convolved with the canonical HRF and its temporal derivatives, to account for potential mismatches in hemodynamic delays (Liao et al., 2002). One then augment the resulting GLM with fMRI confounds (e.g., motion regressors and slow drifts), and fit it to fMRI time series. Fitted regressor weights at each voxel thus provide an estimate 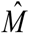 of the local neural response to each trial, which is deconvolved from the HRF and corrected for typical fMRI confounds, and can then enter a mediation analysis. We call this the *deconvolution* approach.

On the other hand, one may reframe the brain-behavior mediation model in the HRF-convolved space. One first resample the explanatory and dependent variables at the fMRI temporal resolution by reweighing each “trial’“ regressor above with its corresponding *X* and *Y* entries and them summing over trials. One then convolves the resulting regressors with the canonical HRF (and its temporal derivatives) and augment the resulting GLM with fMRI confounds prior to entering a mediation analysis. We call this the *convolution* approach.

Both approaches can, in principle, deal with hemodynamic and other fMRI confounds, but they differ in terms of their respective bias-variance tradeoff. The *convolution* approach effectively yields reliable neural response estimates, under the implicit assumption that the HRF is identical across trials. In contrast, the *deconvolution* approach allows for trial-by-trial variations in HRF, at the cost of compromising the reliability of neural response estimates.

In the Results section below, we evaluate the robustness of these two strategies w.r.t. deviations to canonical HRF models.

### A note on causality

Let us now highlight a possible interpretational issue of mediation analysis. Note that Equation 2 implicitly assumes a cascade of causal influences (MacKinnon et al., 2002), which may be best summarized in terms of the directed acyclic graph depicted on Figure 2 below (left panel).

One would then be tempted to interpret a statistically significant mediated effect in causal terms, as in: perturbing the independent variable *X* should result in changes in the mediator variable *M* that would eventually cascade down to the dependent variable *Y*. In the context of brain-behavior mediation, this causal interpretation aligns with the intuitive notion that behavioral responses to stimuli necessarily has to emerge from an intermediate neural information processing step. This causal reasoning, however, does not hold regarding the relationship between *M* and *Y*, which are both *observed* data. In Equation 2, the strength of this relationship is controlled by the path coefficient *b*. Importantly, statistical inference on the path coefficient *b* is but a quantitative assessment of the conditional mutual information *I*(*M*,*Y*|*X*), which is invariant under a reversal of the directionality of the relationship between *M* and *Y*. In other terms, Equation 2 is formally equivalent to the following alternative model:

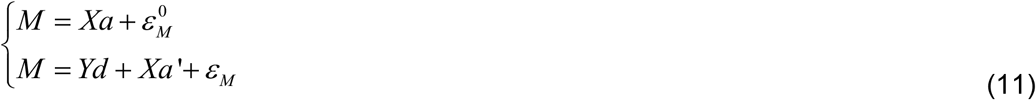

where the second line simply derives from swapping (with impunity) the explanatory and response variables in the second line of Equation 2. Here, *d* and *a′* are “swapped” path coefficients that have a different causal interpretation (cf. Figure 1, right panel), *ε_M_* and are model residuals that are not equivalent to the neural noise 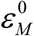 of Equation 2. One can show (see Appendix E) that both native and swapped path coefficients estimates are analytically related as follows:

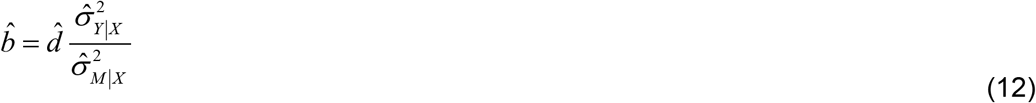

where 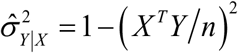 is the sample variance of Equation 1’s residuals estimates 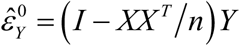. It should be clear from Equations 11–12 that assessing the conditional mutual information *I*(*M*,*Y*|*X*) can be equivalently addressed either by assessing the evidence for *b* ≠ 0 (native form of the mediation model, cf. Equation 2), or by assessing the evidence for *d* ≠ 0 (cf. Equation 11). In fact, the ensuing t-statistics *t_b_* and *t_d_* are exactly equal (see Appendix E), i.e. brain-behavior mediation test statistics are invariant under a permutation of *M* and *Y* variables.

This has two important consequences.

First, one may rely on Equations 11–12 to improve the computational efficiency of brain-behavior mediation analysis by several orders of magnitude. Recall that in the context of whole-brain fMRI, working with regression models where fMRI signals only enter as dependant variables is computationally very advantageous. This is because many algebraic operations that are required for parameter estimation (e.g., here, matrix multiplications and inversions, etc) can be computed once and for all. In brief, the computational gain of performing brain-behavior mediation analysis using Equations 9–10, when compared to Equation 2, is of the order of 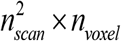, where *n_scan_* and *n_voxel_* are the number of fMRI time samples and voxels, respectively. This may speed up whole-brain mediation analysis by several orders of magnitude.

Second, a statistically significant mediated effect is compatible with two causal interpretations. In particular, under the “swapped” model of Equation 11, variations in behavior *Y* may cause changes in the neural response *M* (cf. Figure 1, right panel). This alternative causal interpretation (*Y → M*) is not as nonsensical as it may first sound. For example, somatosensory cortices will respond to variations in motor actions, eventually enabling proprioceptive sensations. More generally, a given brain system may be collecting and/or processing information regarding overt behavior (which may have been produced elsewhere in the brain) for the purpose of, e.g., learning, memory, metacognition, etc… In any case, this interpretational issue is important, because the implicit intention behind brain-behavior mediation analysis is clearly to provide statistical evidence for the “native” causal scenario (*x →M → Y*). We will comment on this and related issues in the Discussion section of this manuscript.

One may think that affording evidence for the “native” causal claim of brain-behavior mediation analysis may require non observational studies, e.g., causal perturbations of neural activity (lesion studies, transcranial magnetic stimulation, etc). Nevertheless, we argue that one may perform complementary data analyses that may partially address the interpretational issue above. For example, having assessed the significance of a mediated effect, one may exploit locally multivariate information to provide statistical evidence for or against candidate causal claims. In fact, when considering the set of mediator variables within a significant cluster together, “native” (*M → Y*) and “swapped” (*Y → M*) causal interpretations induce a many-to-one and a one-to-many M-Y mapping, respectively (see Figure 3 below). Because “native” and “swapped” causal scenarios differ in terms of whether *Y* is viewed as an input or as an output of local neural information processing, we refer to the ensuing test statistics as an *I/O test statistics*.

**Figure 3:**
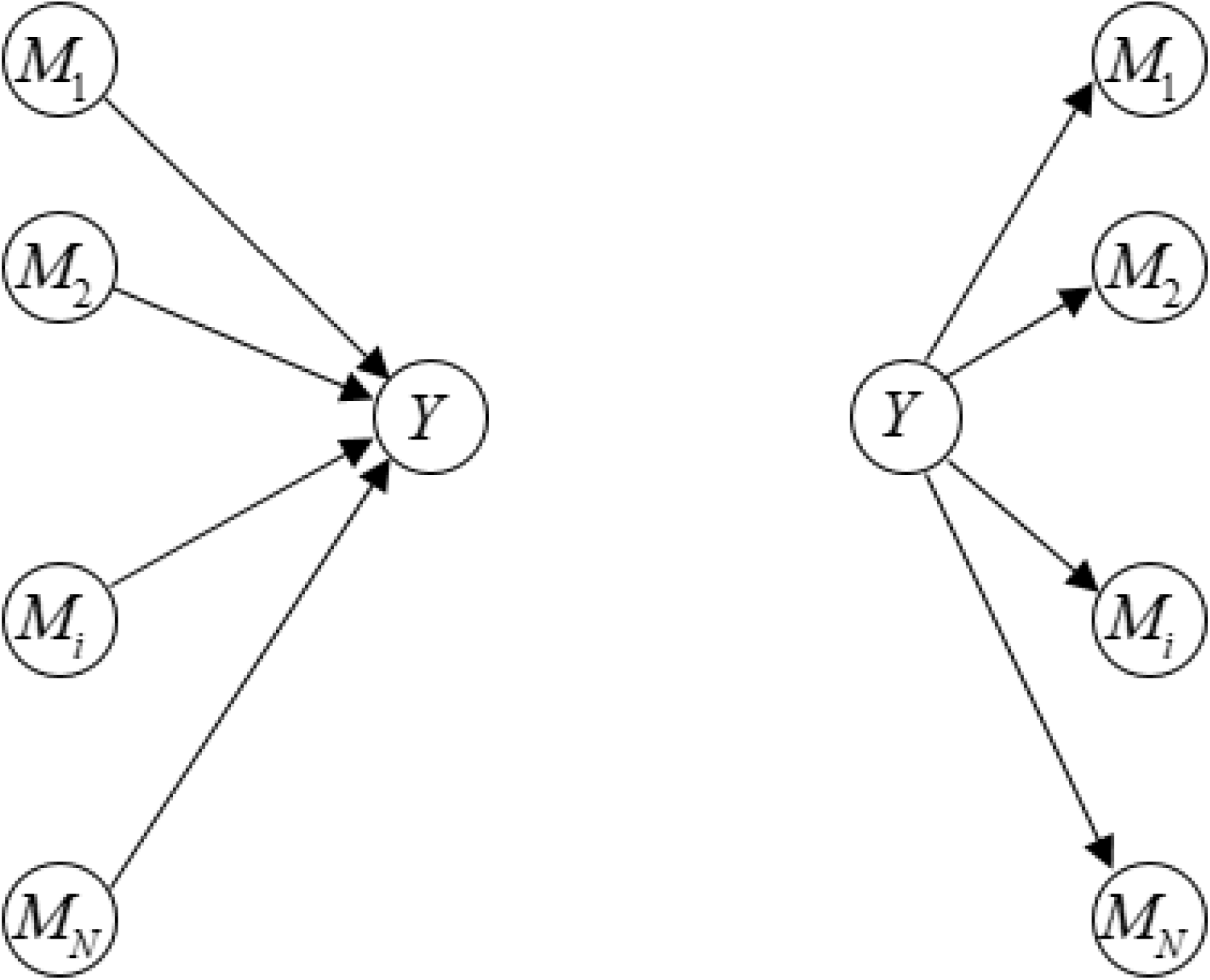
Candidate multivariate input/output M-Y mappings. Left panel: “native” causal interpretation (many-to-one input/output mapping, Y as an output). Right panel: “swapped” causal interpretation (one-to-many input/output mapping, Y as an input).

Let *M_i_* be the trial-by-trial variations of a voxel belonging to a given mediator cluster, where *i* ∈[1, *N*] and *N* is the number of voxels in the cluster. We define our I/O test statistics 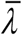 as follows:

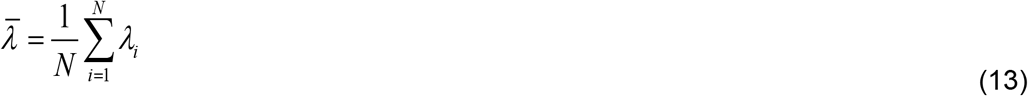

where *λ_i_* is the loss of conditional mutual information between *M_i_* and *Y* when accounting for other neighboring voxels *M_j≠i_*:

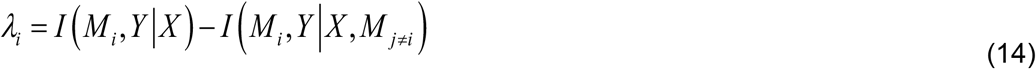

In Equation 14, *I*(*M_i_*, *Y*|*X*, *M_j≠i_*) and *I*(*M_i_*, *Y*|*X*) are the conditional mutual information between *M_i_* and *Y*, given *X* and the activity in all other voxels *j* ≠*i* or not, respectively. Note that *λ_i_* is sometimes coined the *interaction information* (McGill, 1954). In brief, 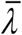 measures the average improvement or worsening of the mutual information between candidate mediator voxels and behavioral responses, when accounting for variations in neighboring brain activity.

For arbitrary gaussian variables *X* and *Y*, the mutual information *I*(*X*,*Y*) can be written as *I*(*X*,*Y*) =−1/2log(1 − *ρ_X,Y_*^2^), where *ρ_X,Y_* is the correlation between *X* and *Y* (Marrelec et al., 2005). In turn, Equation 12 can be rewritten as follows:

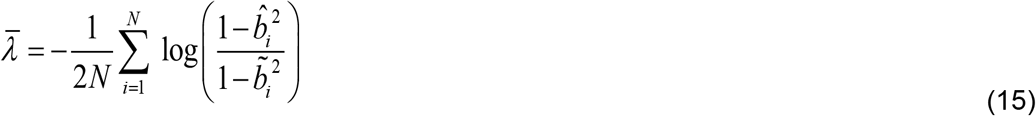

where 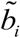 is the conditional correlation between *M_i_* and *Y*, given *X* and *M_j≠i_*:

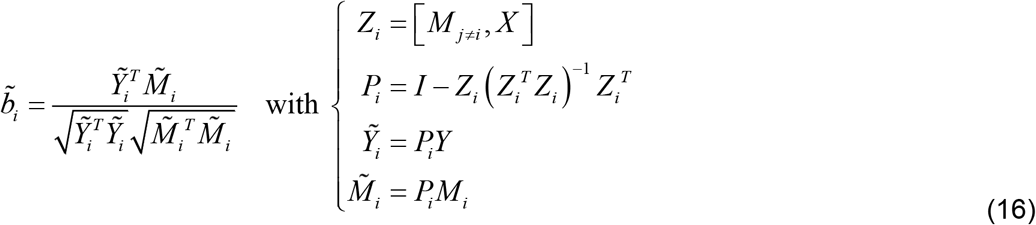

It turns out that the sign of 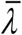 provides evidence in favor or against the native causal interpretation of the brain-behavior mediation model. More precisely: if *M* → *Y*, then 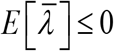, whereas if *Y* → *M*, then 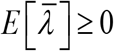. This is because if *Y* is an output of local brain activity (*M → Y*), then any given univariate statistical relationship between *M_i_* and *Y* (path coefficient 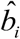) is obscured by the (partially independent) contributions of all other mediator variables *M_j≠i_*. Therefore, when removing all the variability that can be explained with *M_j≠i_*, one reveals the unique contribution of 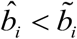. In contrast, if *Y* is an input to local brain activity (*Y → M*), then the variability shared by all mediator variables results from the influence of *Y*. Therefore, when removing all the *M*. variability that can be explained with *M_j≠i_*, one degrades the statistical relationship between *Mi* and *Y* (i.e. 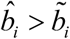). We will evaluate the utility and robustness of our I/O test statistics in the Results section below.

## Results

In what follows, we will be comparing five testing approaches: Sobel’s test, Aorian’s test, Goodman’s test, the M3 bootstrap test, and the conjunctive approach, in terms of their statistical sensitivity and specificity. Using numerical simulations, we will assess the impact of neural noise and deviations to HRF assumptions. Taken together, these *in-silico* experiments will serve to address questions Q1 to Q3. Using further numerical simulations, we will demonstrate the utility of our I/O test statistics for addressing the main interpretational issue of brain-behavior mediation analysis (Q4). Finally, we will report the results of a brain-behavior mediation analysis in the context of an fMRI experiment on decision making under risk.

### Comparing the statistical specificity and sensitivity of testing approaches

First, we ask whether candidate testing approaches yield valid inferences, i.e. whether they allow for a faithful control of false positive rate. To address this question, we simulated data under three different variants of the null hypothesis. More precisely, we simulated 40,000 datasets with Equation 2, using three different settings of the path coefficients, i.e.: (i) *a* = 0 and *b* = 1/2, (ii) *a* = 1/2 and *b* = 0, or (iii) *a* = *b* = 0. In all simulations, we simulated *n* = 50 trials, set the direct effect size to *c′* = 1/2 and used unitary variance for all independent variables in Equation 2 (i.e. *X*, 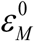 and *ε_Y_*). Across these 40,000 simulations, we then measured the (false positive) detection rate of each candidate testing approach, as one varies the significance threshold *α*. Note that all (indirect or conjunctive) parametric tests were performed with Student’s probability distribution functions with *n* – 2 degrees of freedom. Finally, we kept the default number of 1000 resamplings in the bias-corrected M3 bootstrap test.

Second, we asked how sensitive are candidate testing approaches under moderate mediated effect sizes. Here, we simulated 40,000 datasets with Equation 2, using *a* = *b* = 1/2, and measured the (true positive) detection rate of each candidate testing approach, as one varies the significance threshold *α*.

The results of these analyses are summarized on Figure 4 below.

**Figure 4:**
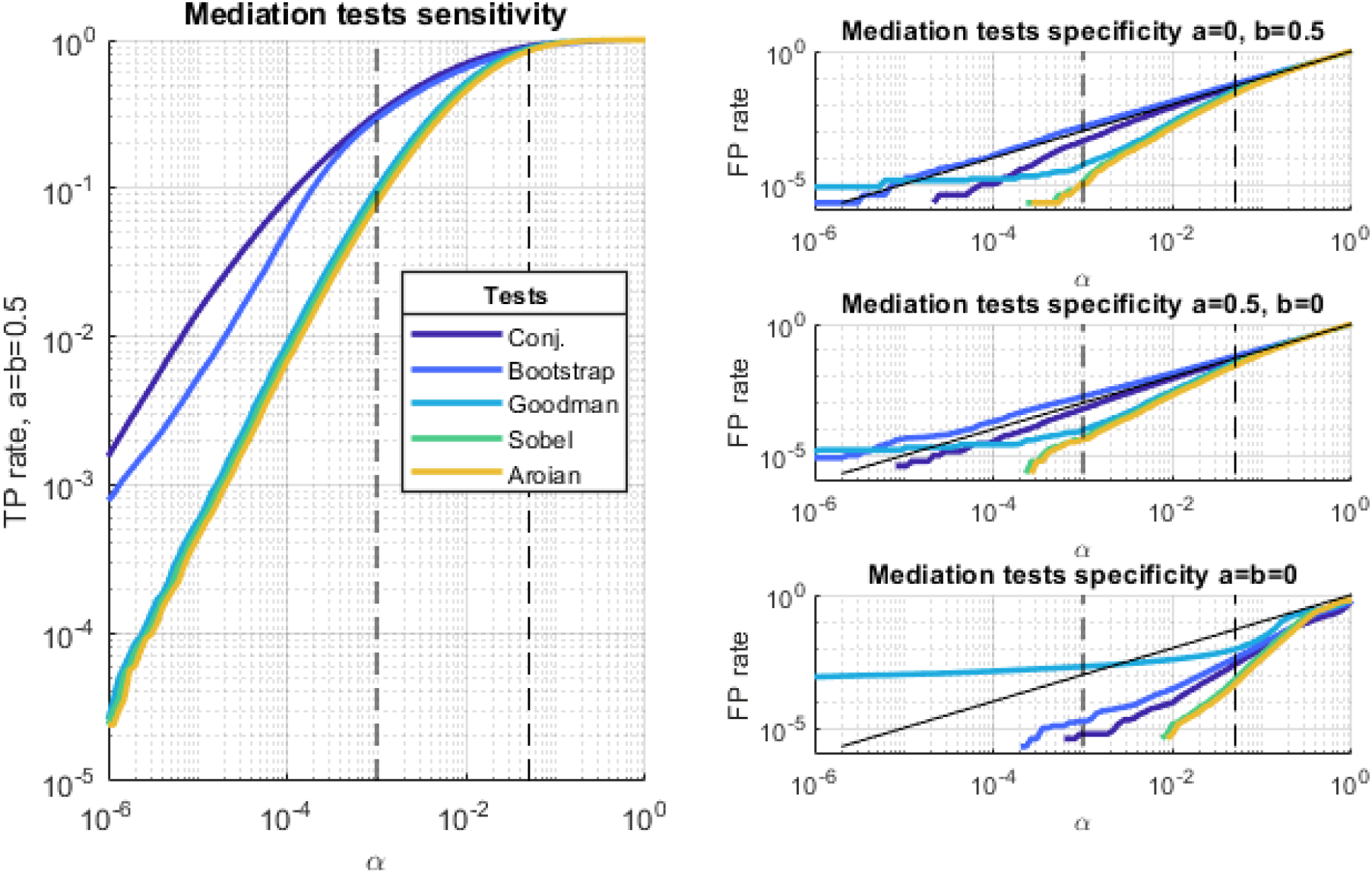
Statistical specificity and sensitivity of variants of mediation significance testing approaches. Left panel: The sensitivity of mediation tests (y-axis) is plotted against the significance threshold α (x-axis), for each candidate testing approach (dark blue: conjunctive testing, blue: M3 bootstrap indirect approach, light blue: Goodman’s indirect approach, green: Sobel’s indirect approach, yellow: Aorian’s indirect approach). Upper right panel: The specificity of mediation tests (y-axis) is plotted against the significance threshold, for H0: a=0 and b=1/2 (same format as left panel). Middle right panel: Same format as upper right panel for H0: a=1/2 and b=0. Lower right panel: Same format as upper right panel for H0: a=b=0.

As expected, the conjunctive test is more sensitive than both Sobel and Aorian tests. Slightly more surprising maybe is the fact that the conjunctive test turns out to also be more sensitive than Goodmans test and the M3 bootstrap test, though the latter reach similar sensitivity levels for significance thresholds higher than 0.001. We will refine our evaluation of statistical sensitivity when assessing the impact of neural noise below.

In addition, all approaches except Goodman and the M3 bootstrap tests are valid, i.e. they yield a false positive rate that is equal or smaller than the significance threshold *α*. Goodman’s test always yield invalid inference if the significance threshold is small enough, whereas the M3 bootstrap test only yields invalid inference when *b* = 0. Note that the conjunctive approach is the least conservative of all tests, and this difference grows when the significance threshold decreases.

### Assessing the impact of neural noise

Recall that the magnitude of neural noise is expected to play a critical role for the statistical sensitivity of mediation analysis. To demonstrate this effect, we simulated 10,000 datasets using the same parameter settings as above, except for neural noise magnitude, which we varied from 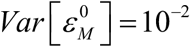 to 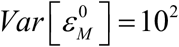. For each neural noise magnitude, we then measured the (true positive) detection rate of each candidate testing approach, when setting the significance threshold to *α* = 0.005. The ensuing sensitivity profiles are summarized on Figure 5 below (left panel).

**Figure 5:**
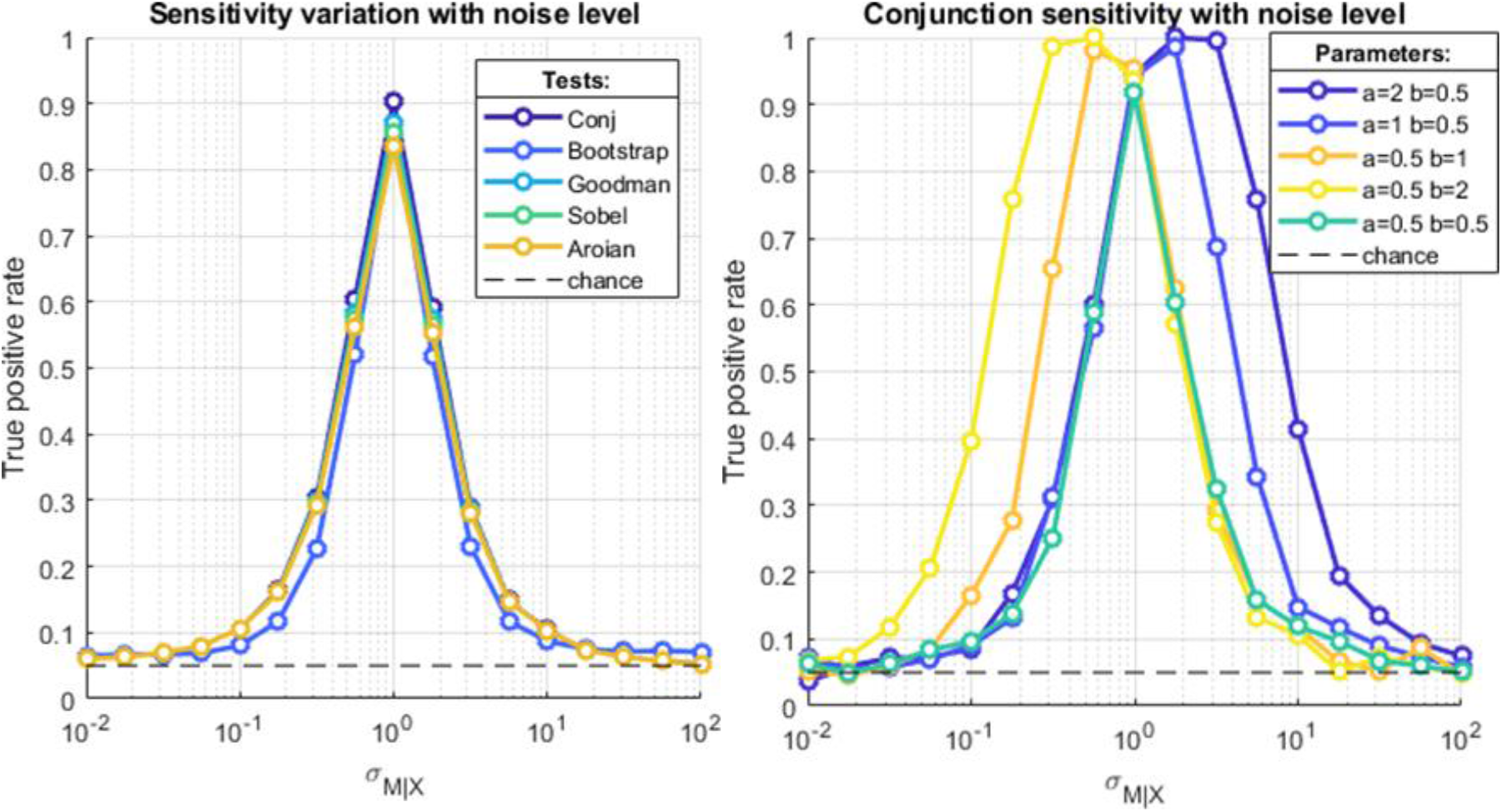
The impact of neural noise on statistical power. Left panel: The sensitivity of mediation tests (y-axis) is plotted against the variance of neural noise (x-axis), for each candidate testing approach (same format as Figure 4), when a=b=1/2. Chance level is indicted using a black dotted line. Right panel: The sensitivity of conjunctive mediation tests (y-axis) is plotted against the variance of neural noise (x-axis), when varying path coefficients (dark blue: a=2 and b=1/2, blue: a=1 and b=1/2, cyan: a=b=1/2, orange: a=1/2 and b=1, yellow: a=1/2 and b=2).

All testing approaches have a similar sensitivity profile, which follows a bell-shaped function of neural noise magnitude, with an apex around 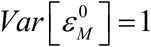. This corresponds to a situation in which about 20% of the trial-by-trial variance in *M* is explained by *X*. As the amount of explained variance in *M* departs from this nominal level, the sensitivity of mediation analysis effectively tends towards chance level. Now, everything else being equal, increasing *a* or the variance of *X* eventually inflates sensitivity on the right tail of the sensitivity profile, while increasing *b* rather boosts its left tail. This moves the position of sensitivity apex towards smaller and stronger noise variance, respectively (see Figure 5, right panel).

Now, in the Methods section above, we reasoned that the expected sensitivity profile of mediation analysis should eventually favor the detection of neural information processing steps that are positioned away from either end of the processing hierarchy. In what follows, we compare candidate testing approaches w.r.t. their ability to detect levels in a simple feed-forward hierarchy. In brief, we simulated 1,000 datasets under Equation 10, using 100 intermediary network nodes. In all simulations, initial and final path coefficients were set to *a*_0_ = *b* = 1/2 and all intermediary path coefficients were set to *a* = 1 ∀*i*. In addition, the variance of all independent variables were set to unity except for the local neural noise increments, whose standard deviation was set to 0.3. Following the principle of mass-univariate mediation analysis, a mediation test was then performed on each node in isolation (significance threshold: *α* = 0.005). For each network node, the ensuing (true positive) detection rate was then measured across the 1,000 simulations. The result of the ensuing detection profile is shown on Figure 6 below.

**Figure 6:**
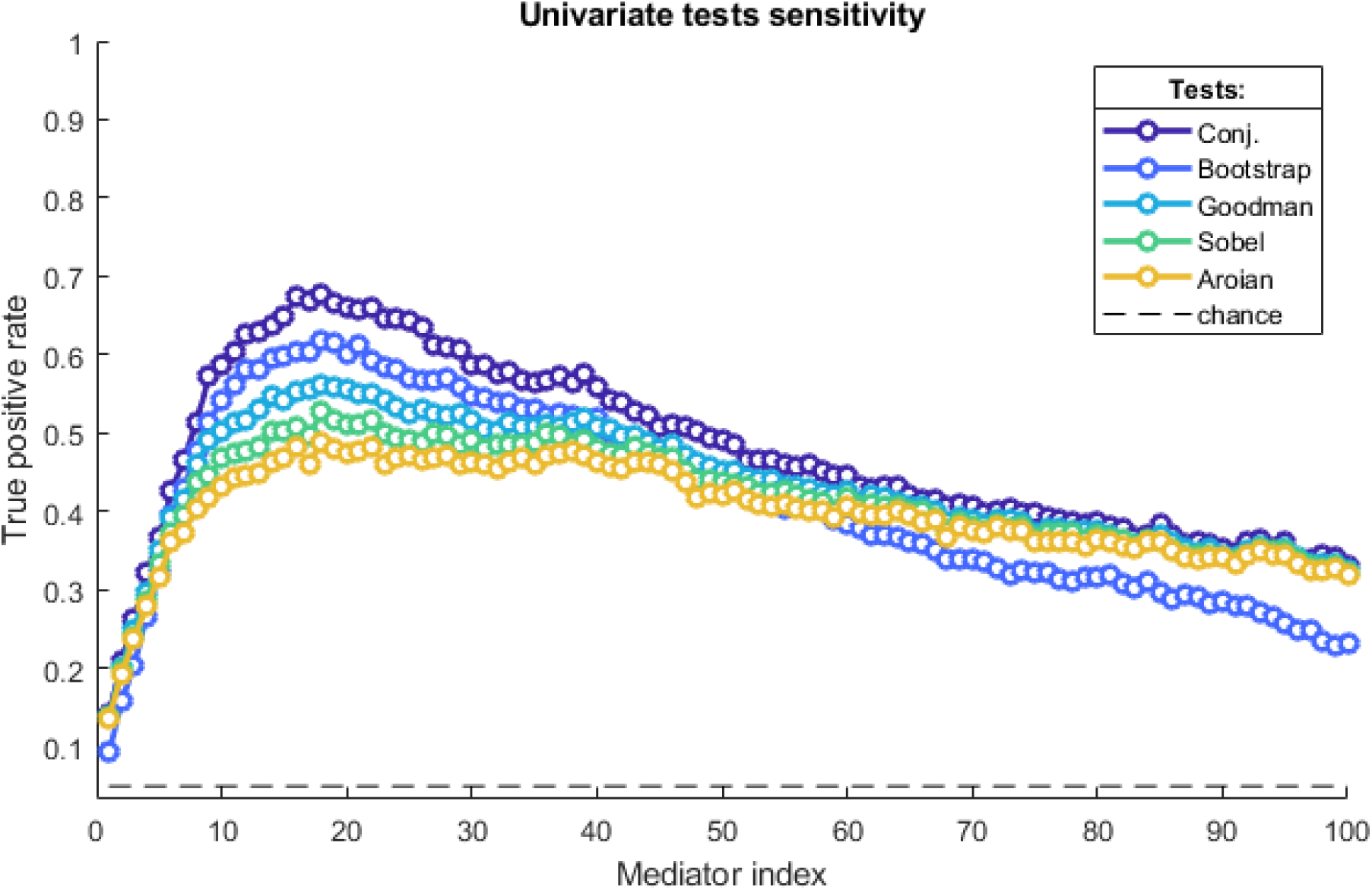
The heterogeneity of statistical sensitivity. The sensitivity of mediation tests (y-axis) is plotted against the hierarchical level of candidate mediators along the serial processing pathway (x-axis), for each candidate testing approach (same format as Figure 4).

As expected, local neural noise increments accumulate along the hierarchy, effectively increasing the neural noise level estimate as the hierarchical level increases. In turn, the detection profile also follows a bell-shaped function of hierarchical level, such that lower and higher hierarchical levels are less easy to detect. Interestingly, one can also see that different testing approaches have different sensitivity profiles. In particular, one can see that the conjunctive approach exhibits a higher sensitivity than all other approaches, irrespective of the hierarchical level of interest. Note that the M3 bootstrap test is better than other indirect approaches for intermediate hierarchical levels, but eventually loses its competitive advantage for higher hierarchical levels.

### Assessing the robustness to deviations from hemodynamic assumptions

Despite the inclusion of HRF derivatives in the mediation model, deviations to the canonical HRF can impair test sensitivity. In this section, we compare the robustness of *convolution* and *deconvolution* approaches to unanticipated delays in HRF. We thus simulated 100 datasets using the same parameter settings as above, except that we varied systematically the HRF delay, effectively inducing a shift with the canonical HRF ranging from −5 second to 5 seconds. Each dataset was then analyzed using all (*indirect* and *conjunctive*) testing approaches, under both *convolution* and *deconvolution* strategies (with the canonical HRF and its delay derivative). For each HRF delay shift, we then measured the (true positive) detection rate of each candidate mediation analysis strategy, when setting the significance threshold to *α* = 0.005. The ensuing sensitivity profiles are summarized on Figure 7 below.

**Figure 7:**
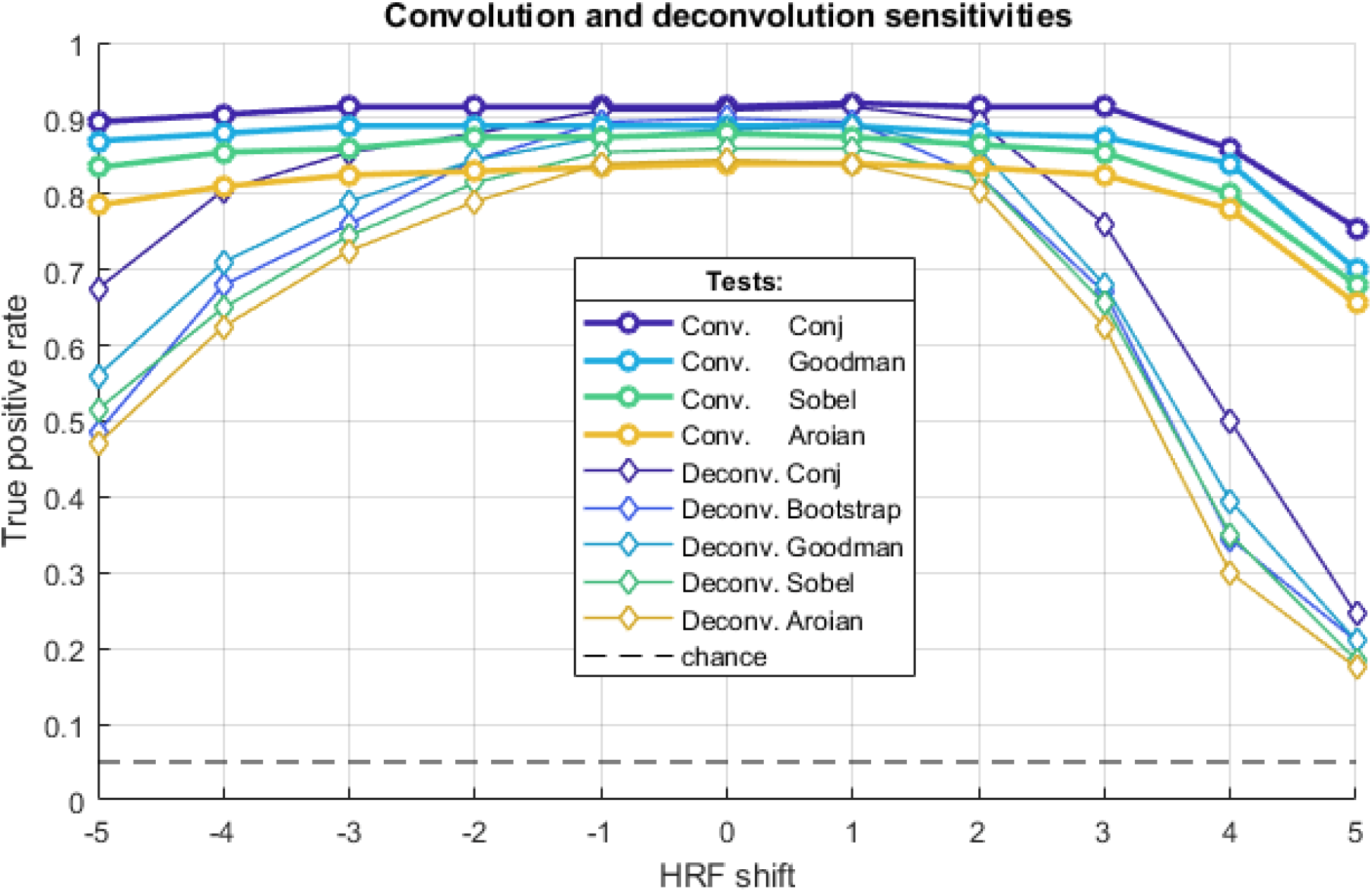
The impact of unmodelled hemodynamic delays. The sensitivity of mediation tests (y-axis) is plotted against the HRF shift (x-axis), for each candidate testing approach (same format as Figure 4, thick lines: *convolution* approach, thin lines: *deconvolution* approaches).

All mediation analysis strategies exhibit a bell-shaped sensitivity profile, eventually peaking when there is no deviation to the canonical HRF (i.e. when the HRF delay shift is null). Also, when there is no deviation to the canonical HRF, *deconvolution* and *convolution* strategies yield similar test sensitivity. However, when the deviation to the canonical HRF increases, the loss of statistical sensitivity is much stronger for *deconvolution* than for *convolution* approaches. For example, with a (realistic) delay shift of 3 seconds, most *deconvolution* approaches lose about 10% to 15% sensitivity on average. In comparison, *convolution* approaches only lose about 2% sensitivity. In addition, the *conjunctive* approach always exhibit higher sensitivity than *indirect* approaches, irrespective of whether one chooses a *convolution* or *deconvolution* strategy.

We note that, with a significance threshold of *α* = 0.05, deviations to the canonical HRF has no adverse effect on the validity of statistical tests, i.e. all mediation test approaches yield 5% or less false positive rates under the null.

### Addressing the interpretational issue of brain-behavior mediation analysis with the I/O test statistics

Recall that a significant mediated effect may have two distinct causal interpretations: the behavioral variable may either be an input (*Y → M*) or an output (*M → Y*) of the brain region where the null has been rejected. To address this issue, we proposed a simple I/O test statistics, whose sign is expected to discriminate between these two scenarios.

Here, we evaluate the utility of the I/O test statistics 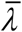, in conditions similar to our fMRI data analysis below, using numerical Monte-Carlo simulations.

First, we simulated data under three scenarios:

- H_1_ (native causal scenario *M →Y*): the independent variable *X* is sampled under a normal distribution, each multivariate mediator unit *M_i_* is set to a noisy affine transformation of *X* (with random weights), and the dependent variable *Y* is set as a noisy mixture of *X* and all mediator units (with random weights).
- H_2_ (“swapped” causal scenario *Y →M*): the independent variable *X* is sampled under a normal distribution, the dependent variable *Y* is set as a noisy affine transformation of *X* (with a random weight) and each multivariate mediator unit *M_i_* is set to a noisy mixture of *X* and *Y* (with random weights).
- H_0_ (null scenario): the independent variable *X* is sampled under a normal distribution, and all other variables are set to a noisy affine transformation of *X* (with random weights).

We simulated each scenario 1000 times, with 64 trials and 20 mediating units (all random variables and weights were sampled under a centered normal distribution with unit variance). Note that, in all three scenarios, *M* and *Y* variables are correlated with each other (under the null, this is because of the influence of *X*, which acts as a confounding variable). For each simulated dataset, we derive the I/O test statistics 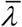. The resulting Monte-Carlo distributions are shown on Figure 8 below (left panel).

**Figure 8:**
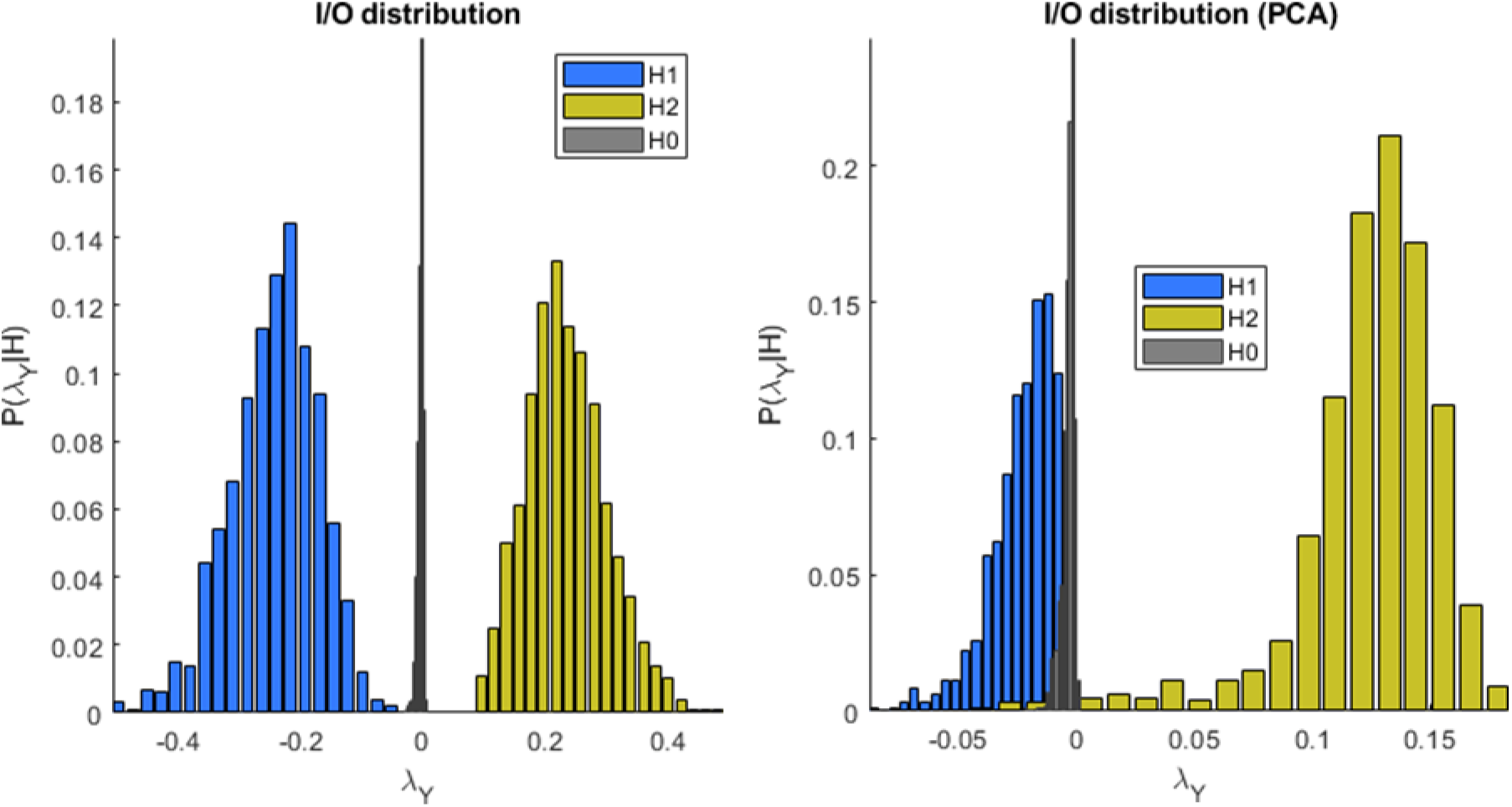
Sensitivity and robustness of the I/O test statistics λ. Left panel: The Monte-Carlo distribution of the I/O test statistics λ (y-axis) is plotted under alternative scenarios (H1: blue, H2: yellow, H0: grey). Right panel: Same format as left panel, but under data dimension reduction (20 first principal components of a PCA).

On can see that the three scenarios are very well discriminated. In particular, the distribution of the 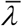 under the null is centered on zero, and lies in between its distribution under H_1_ and under H_2_. Moreover, and as expected, 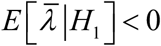 and 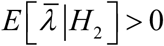.

These simulations however, do not account for the limitations that arise in realistic settings. In particular, the number of neural units or voxels that compose the multivariate set of mediators may largely exceed the number of trials. Here, a pragmatic solution is to perform a PCA decomposition, and keep the *K* first principal components to summarize the within-region variability. We now ask whether the ensuing I/O test statistic is robust to this dimension reduction. In brief, we performed the same set of simulations as above, this time simulating 100 mediating units and deriving the test statistics from the ensuing *K*=20 first principal components. The resulting Monte-Carlo distributions are shown on the right panel of Figure 8.

One can see that the dimension reduction strongly reduces the range of variation of the I/O test statistics, when compared to the situation above, where all the relevant variation is available. Furthermore, the magnitude of 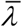 under scenario H_1_ and H_2_ is asymmetrical. More precisely, one can see that 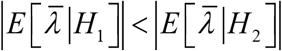. In other terms, when relevant information is lacking, the average evidence in favor of H1 is weaker than the average evidence in favor of H2. Nevertheless, the sign of the I/O test statistics can still be interpreted as evidence for or against the native causal interpretation of the brain-behavior mediation model, i.e. 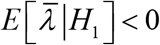 and 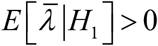.

### fMRI study of decision making under risk

Here, we perform a brain-behavior mediation analysis of previously acquired fMRI data (Chen, 2014), which is openly available as part of the OpenFMRI project (Poldrack et al., 2013). In this study, 60 participants made a series of 64 accept/reject decisions on risky gambles. On each trial, a gamble was presented, entailing a 50/50 chance of gaining an amount G of money or losing an amount L (so-called “baseline” condition). Participants were told that, at the end of the experiment, four trials would be selected at random: for those trials in which they had accepted the corresponding gamble, the outcome would be decided with a coin toss, and for the other ones -if any-, the gamble would not be played. All 64 possible combinations of G/L pairs (10$<G<40$, 5$<L<20$) were presented across trials, which were separated by 7 seconds on average (min 6, max 10). MRI scanning was performed on a 3T Siemens Prisma scanner. High-resolution T1w structural images were acquired using a magnetization prepared rapid gradient echo (MPRAGE) pulse sequence with the following parameters: TR = 2530 ms, TE = 2.99 ms, FA = 7, FOV = 224 × 224 mm, resolution = 1 × 1 × 1 mm. Whole-brain fMRI data were acquired using echo-planar imaging with multi-band acceleration factor of 4 and parallel imaging factor (iPAT) of 2, TR = 1000 ms, TE = 30 ms, flip angle = 68 degrees, in-plane resolution of 2X2 mm 30 degrees of the anterior commissure-posterior commissure line to reduce the frontal signal dropout, with a slice thickness of 2 mm and a gap of 0.4 mm between slices to cover the entire brain. See https://openneuro.org/datasets/ds000053/versions/00001 for more details.

Data preprocessing included standard realignment and movement correction steps. Note that we excluded 2 participants, either due to missing information or because the misalignment between functional and anatomical scans could not be corrected.

We first regressed, for each participant, the observed choices against gains and losses (Equation 1). This yielded estimates of the total effects 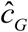 and 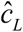 of gains and losses, respectively. This also provided an estimate 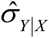 of the behavioral residuals’ standard deviation. The results of this analysis are shown on Figure 9 below.

**Figure 9:**
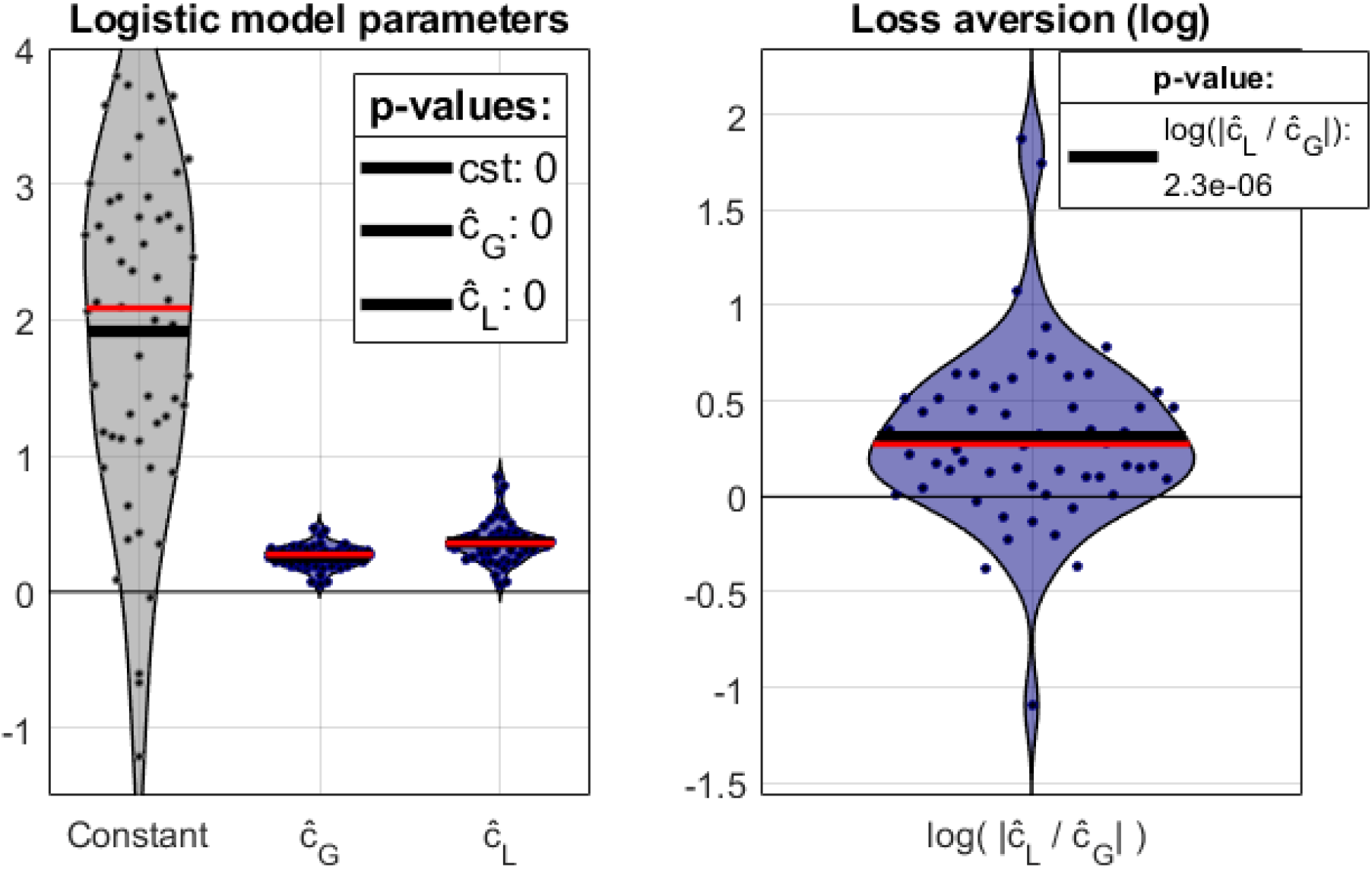
Summary of behavioral results. Left panel: Between-subject empirical distribution of estimated within-subject parameters (left: constant term in the regression, middle: gain weight c_G_, right: loss weight c_L_). The black and red lines show the group-level mean and median, respectively. Right panel: Between-subject empirical distribution of the loss-version index (same format as left panel).

In brief, both gain and loss factors have a significant effect on decisions under risk (gain factor: p<10-5, loss factor: p<10-5). We note that, together, gain and loss factors explain on average 44.6% (std: 24.2%) of the trial-by-trial variance on participants’ decisions (average balanced accuracy: 84.84%, std: 9%).

For each participant, we also derived a loss-aversion index: 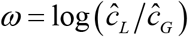, which is positive when losses have a stronger weight on accept/reject decisions than gains. One can see that the average loss-aversion index is significant (p<10-5), i.e. losses have more weight on participants’ decisions than gains.

Then, we analyzed fMRI time series (at the within-subject level) using both *convolution* and *deconvolution* approaches.

The *convolution* strategy relied upon the following two GLMs:

- **Equation 10 (first line):** GLM1 included regressors for trial-by-trial gains and losses (temporally aligned with the gamble presentation and convolved with the canonical HRF and its delay derivative), and basic confounding factors (six movement regressors and their squared values, as well as a Fourier basis set for slow drift removal). Fitting GLM1 to each fMRI voxel time series yielded a map of estimates 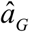 and 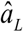 that correspond to the local effect of gain and loss on neural activity at the time of gamble presentation, respectively. In addition, we extracted the standard deviation of GLM1’s residuals, which form a map of the local neural noise’s strength 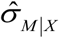.
- **Equation 11 (second line):** GLM2 is identical to GLM1, but also includes acceptance/rejection choices (convolved with the canonical HRF and its temporal derivatives). Fitting GLM2 to fMRI time series yielded regressor weight estimates that measure the correlation between local neural activity and behavior, above and beyond the effect of gain and losses 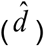. The map of local path coefficients 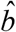 was then obtained from 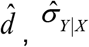 and 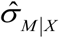 using Equation 12.

The *deconvolution* strategy was implemented as follows. First, we fitted GLM3, which included “trial” regressors (temporally aligned with the gamble presentation) as well as basic fMRI confounds. Regression weight estimates yielded local trial-by-trial neural responses 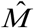. Maps of path coefficients estimates 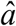 and 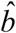 were obtained using Equation 4, given local neural responses 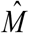.

Random-effect group-level inference on the mediation of gain and loss factors was then performed by reporting group averages of path coefficients 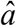 and 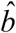, after 8mm FWHM smoothing. We applied all indirect and conjunctive approaches except for the M3 bootstrap method (because of its limited statistical gain, when compared to its computational cost). For all approaches, we used unsigned (two-tailed) tests with standard random field theory (RFT) correction for whole-brain multiple comparisons correction.

In brief, no mediation testing approach based upon the *deconvolution* strategy reached statistical significance. This was the case even when using more lenient corrections for multiple comparisons (e.g., FDR). This was however not the case for mediation analyses based upon the *convolution* strategy. Here, *indirect* approaches yielded group-level significant clusters at low-set inducing thresholds (p=0.01 or p=0.05, uncorrected). In what follows, we discard these results as these thresholds are known to violate RFT assumptions (Flandin and Friston, 2019). Now, under the default set-inducing threshold (p=0.001, uncorrected), the conjunctive approach identified 6 clusters that significantly mediate the effect of gain: the right supramarginal gyrus or SMG (p=0.011, RFT-corrected), bilateral posterior dorsomedial PFC or BA8 (left: p=0.003, right: p=0.008, RFT-corrected), the right anterior ventrolateral PFC or BA45 (p=0.018, RFT-corrected) and bilateral posterior dorsolateral PFC or BA8/9 (left: p=0.007, right: p=0.009, RFT-corrected). In addition, there was a trend (p=0.06, RFT-corrected) for 1 cluster mediating the effect of loss, in the left anterior ventrolateral PFC. These clusters are shown on Figure 10 below.

**Figure 10:**
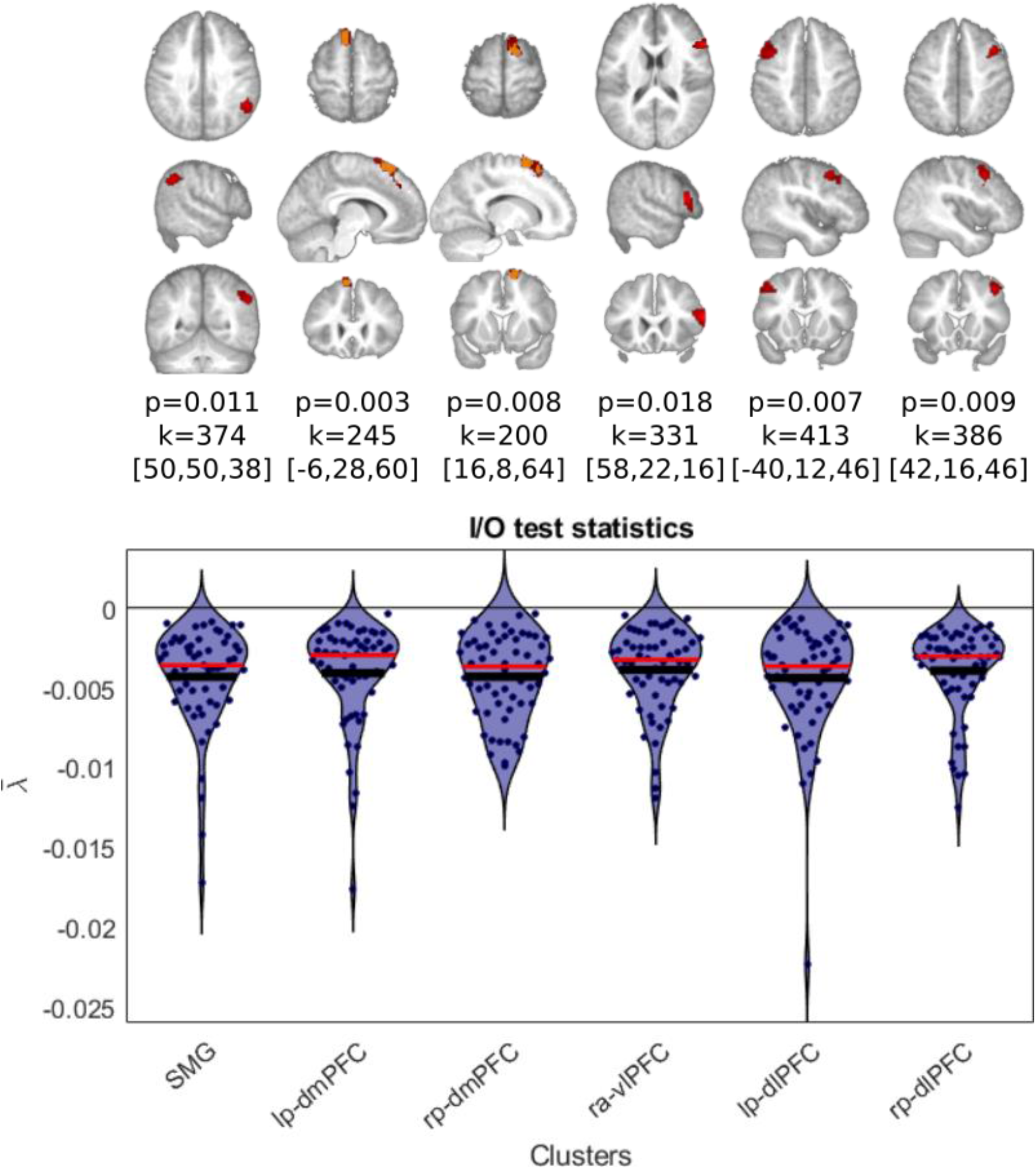
Significant mediators of gains and losses on decisions under risk. Upper panel: the six significant clusters of brain-behavior mediation analysis (*conjunctive/convolution* approach) are shown on axial (up), sagittal (middle) and coronal views (bottom). All maps used a default set-inducing threshold of correction p=0.001 uncorrected (red areas) for the RFT correction, except the bilateral dmPFC’s map where with p=0.0002 uncorrected (yellow areas) in order to separate the two hemispheres. Lower panel: The ensuing between-subject empirical distribution of the I/O test statistics λ (y-axis, group-level mean ±standard deviation) is shown for each significant clusters (x-axis).

We note that regions contralateral to significant unilateral mediators of the gain effect were all close to statistical significance: c.f. left SMG (p=0.094, RFT-corrected) and left anterior vlPFC or BA45 (p=0.177, RFT-corrected).

At the very least, these analyses demonstrate the superior statistical efficiency of *conjunctive/convolution* approaches. In brief, no other candidate variant of mediation analysis yields positive results on this dataset.

Now, the significant mediated effects above may have two distinct causal interpretations. To afford evidence in favor or against the “native” causal claim of brain-behavior mediation analysis, we derived, for each participant and each significant cluster, our I/O test statistics. Note that, prior to the analysis, we summarized the trial-by-trial variance in each cluster using the 20 first principal components from the within-cluster PCA decomposition (on average cross clusters and participants, these preserve 89% ±2% of the trial-by-trial variance). The group-level empirical distribution of 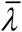 is shown on the lower panel of Figure 10, for each of the 6 significant clusters. Reassuringly, all clusters exhibit strong evidence in favor of the “native” causal interpretation of brain-behavior mediation analysis, i.e. 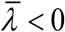 for all subjects and all clusters. We will comment on these results in the Discussion section.

Now, the effect of experimental factors seems to be mediated by a set of anatomically segregated regions in the brain. These regions are likely to be organized into a functional network (cf. Figure 1 above), eventually exerting competitive and/or cooperative influences on behavioral responses. The analysis above is agnostic about the functional architecture of this network. However, the extent to which each of these network nodes actually mediates the effect of gains and losses onto choices varies across subjects. Thus, a given individual may have an idiosyncratic structure of brain pathways for processing gain and loss information. In turn, inter-individual differences in the pattern of mediated effect sizes may have behavioral consequences in terms of how strongly gains and/or losses impact decisions under risk.

Recall that the balance between the behavioral effects of gains and losses is measured using the loss aversion index 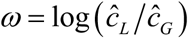 (cf. Figure 9). One may thus ask whether the pattern of mediated effect sizes predicts loss aversion. We thus extracted, in each voxel of the 6 significant mediating clusters above, the indirect effect size 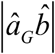, and average these within each cluster. This resulted in 6 region-specific indirect sizes per participant. We then regressed loss aversion indices against (log-transformed) indirect effect sizes, across participants. The results of this analysis are summarized on Figure 11 below.

**Figure 11:**
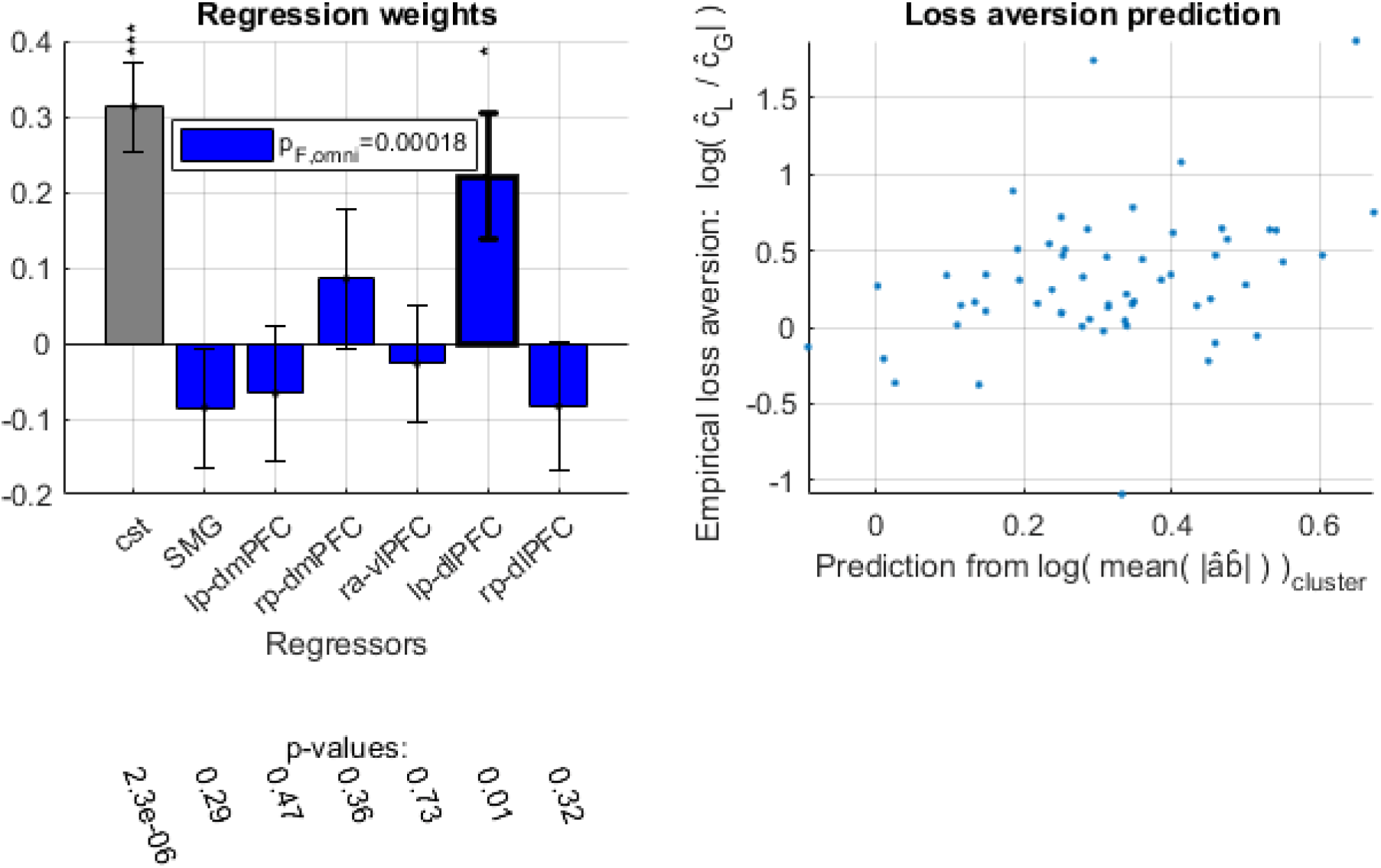
Inter-individual differences in loss-aversion. Left panel: regression coefficients of the analysis of inter-subject differences of loss aversion (grey: constant term, blue: weight of inter-individual differences in cluster-averages of indirect effect size). Errorbars depict standard errors of the mean. Right panel: Observed (y-axis) versus predicted (x-axis) loss aversion. Each dot is a participant.

First, an omnibus F-test shows that the pattern of indirect effect sizes significantly predicts loss aversion (F=5.13, dof=[7,51], R2=12.8%, p=2×10^-4^). This is important, since this means that one can think of loss aversion in terms of a trait that is partly determined by the relative contribution of processing pathways that mediate the effect of gains onto decisions under risk. In addition, one can see that loss aversion increases when the indirect effect size in the left posterior dmPFC increases (t=2.66, dof=51, p=0.01). No indirect effect size in any other region has a significant effect on loss aversion (all p>0.29).

## Discussion

In this work, we identified the specific challenges of brain-behavior mediation analysis. In particular, we evaluated the specificity and sensitivity of five statistical tests, including so-called *indirect* and *conjunctive* approach. In brief, the *conjunctive* approach systematically shows higher sensitivity, while yielding valid inference. In addition, we disclosed the non-trivial impact of neural noise, and assessed the robustness to deviations from fMRI modelling assumptions. The former implies that brain-behavior mediation analysis cannot detect mediators that are too close from either end of the neural information processing hierarchy. *In-silico* investigations of the latter eventually favor the *convolution* approach to HRF modelling. We also disclosed some interpretational issues of mediated effects, in particular: significant mediated effects have two distinct causal interpretations. Importantly, this causal degeneracy may be partially addressed using complementary multivariate I/O test statistics. In addition, it has unexpected favorable computational consequences for whole-brain mediation analysis. Lastly, brain-behavior mediation analysis of fMRI data acquired in the context of decisions under risk further demonstrated the importance of methodological choices regarding brain-behavior mediation analysis. Eventually, the *conjunctive*/*convolution* test approach showed that the right SMG, bilateral posterior dmPFC, right anterior vlPFC and bilateral posterior dlPFC mediate the effect of prospective gains on decisions under risk. Group-level I/O test statistics provided evidence that these regions are contributing to shaping behavioral responses (in a feedforward, causal, manner), rather than collecting and/or processing information about it (cf. interpretational issue). Finally, we showed that inter-individual differences in loss aversion is partly determined by the relative contribution of these six regions to behavioral control.

Taken together, our numerical simulations and analyses of experimental fMRI data demonstrated that *conjunctive* testing has higher statistical sensitivity than *indirect* approaches. This is true even for the bias-corrected M3 bootstrap test, despite its huge computational cost. We note that the sensitivity of the M3 bootstrap test may, in principle, be improved by increasing the number of permutations used to approximate the null distribution (here: 1,000). This however, would render whole-brain analysis excessively slow. Note that M3 bootstrap and conjunctive tests had already been compared at the standard 5% significance threshold outside the context of fMRI (Hayes and Scharkow, 2013). Although authors noted that bias-corrected bootstrap tests were slightly invalid (false positive rate greater than 5%), they recommended them because they eventually yielded more reliable confidence interval estimations. We extended these simulations, eventually showing that the invalidity of bias-corrected bootstrap tests increases as one relies on more stringent significance threshold (cf. Figure 3), which is required when correcting for multiple comparisons. For all these reasons (test validity, statistical sensitivity and computational cost), we would rather favor conjunctive testing for mass-univariate brain-behavior mediation analysis.

Although computationally expedient, mass-univariate brain-behavior mediation analysis essentially relies upon an incomplete model. Not only is it agnostic about the structure of the distributed brain system that process the incoming information (cf. Figure 1), but local, voxel-based, mediation tests simply ignore about 99.999% of the brain. We would argue however, that such incompleteness may be *necessary* for statistical mediation analysis. Recall that evidence for a mediated effect requires an appropriate amount of neural noise. But neural noise estimates have two entirely distinct sources. On the one hand, it may derive from irreducible variations in neural responses that are inherent to the underlying neurobiological processes. On the other hand, it may arise from imperfections in the way neural responses are modeled. The latter most likely applies to the linear brain-behavior model in Equation 2. For example, saturating neural responses to stimuli would, under Equation 2, inflate model residuals. However, although the ensuing neural noise estimates 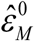 would be partly artefactual, they would still be very informative to predict behavioral responses *Y* above and beyond the *linear* effect of *X*. Now, let us assume that a neurocognitive model was available, that would describe how incoming information *X* would be distorted, transformed and integrated with other (potentially incidental) processes, along the processing hierarchy. For example, such model may derive from recent work in theoretical neuroscience regarding population coding (Averbeck et al., 2006; Georgopoulos et al., 1986), predictive coding (Bastos et al., 2012; Hosoya et al., 2005; Rao and Ballard, 1999) or efficient coding (Barlow, 1961; Doi and Lewicki, 2011). Or it could rely on agnostic multivariate and/or nonlinear decompositions that, when properly parameterized, would account for all sorts of complex relationships between *X* and *M*. In any case, if the model was complete enough, then observed neural activity would not strongly deviate from its predictions. This would preclude the statistical detection of mediated effects. Ironically speaking then, progress in modelling neural information processing may eventually hinder the statistical efficiency of brain-behavior mediation analysis. More practically, this means that statistical brain-mediation analysis may be used in an exploratory manner, to identify brain regions that contribute to behavioral control. Further, complementary, model-based approaches to neural information processing would then help reducing one’s epistemic uncertainty regarding neural noise. For example, artificial neural network modelling may be useful to identify either the structure of processing pathways (Rigoux and Daunizeau, 2015) and/or the impact of incidental biological constraints that may distort local neural information processing (Brochard and Daunizeau, 2020).

This is not to say, however, that statistical brain-behavior mediation analysis cannot be improved.

For example, one may aim at providing more informative inferences regarding the structure of the underlying processing hierarchy. A possibility here is to merge mediation analysis with existing graph analysis techniques that were developed for assessing effective connectivity in the brain (Alstott et al., 2009; Smith et al., 2011; Sporns, 2013). Another, less exhaustive but simpler, solution is to work iteratively: having identified a brain region that significantly mediates the *X* → *Y* effect, one may then look for other brain regions that would mediate both *X* → *M* and *M* → *Y* relationships, and repeat on subsequent mediators. Note that this would require additional corrections for the natural dependencies between brain regions. We refer the interested reader to Van Kesteren & Oberski (2019) for an interesting first step in this direction.

We also think that progress can be made regarding the main interpretational issue of brain-mediation analysis. In this context, let us highlight two extensions of linear mass-univariate approaches that sound promising.

First, one may rely on more stringent inferences regarding the causality of the *M* → *Y* relationship (Preacher, 2015). For example, temporal precedence may be accounted for, and inserted in the brain-mediation model using variants of Granger causality (Zhao and Luo, 2017). Note that special care must be taken regarding hemodynamic delays, whose variations across brain regions may confound temporal precedence. In particular, established fMRI applications of Granger causality are known to be prone to such confounds (David et al., 2008; Deshpande et al., 2010; Zhao and Luo, 2017). Nevertheless, constraining the brain-behavior mediation model with temporal precedence would likely reduce spurious inferences.

Second, one may exploit locally multivariate information to discriminate between many-to-one (*M* → *Y*) and one-to-many (*Y* → *M*) input/output mappings. In this work, we proposed a first step in this direction: namely, the I/O test statistics 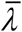. Numerical simulations demonstrated the utility of this information-theoretic measure, and its robustness to partial information losses that result from data dimensionality reduction. However, this work falls short of an exhaustive analytical treatment of I/O test statistics. For example, neither did we investigate whether and how nonlinearities in causal relationships confound and/or bias 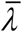 estimates, nor did we derive a formal statistical test of the significance of 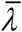 estimates. We note that the difficulty here, is that the null hypothesis may not be the most useful reference point for I/O test statistics. Rather, one aims at comparing two alternative non-nested models. Therefore, an optimal statistical treatment of I/O tests statistics may be best approached using a bayesian approach (Kass and Raftery, 1995; Liu and Aitkin, 2008). We will pursue this and related extensions of I/O test statistics in subsequent publications.

Finally, let us discuss the results of our fMRI analysis. Recall that we identified six candidate mediators of the effect of gain onto decisions under risk. Among these, the posterior dmPFC was previously shown to regulate speed-accuracy tradeoffs (Forstmann et al., 2008) and its anatomical lesion is known to impair inhibitory control in the presence of response conflict (Nachev et al., 2007). Also, decades of neuroimaging, stimulation and lesions studies have evidenced the role of posterior dlPFC and vlPFC cortices in cognitive control (Gbadeyan et al., 2016; Levy and Wagner, 2011; MacDonald et al., 2000; Miller and Cohen, 2001; Nee and D’Esposito, 2017; Soutschek and Tobler, 2020). In addition, functional and anatomical studies report convergent evidence that the right SMG is crucial for regulating emotional responses (Adolphs, 2002; Makovac et al., 2016;

Silani et al., 2013). Now, in the context of decisions under risk, automatic fear responses may induce a default tendency to reject risky gambles, eventually yielding loss aversion (Martino et al., 2006, 2010). This default emotional bias may enter in conflict with the appetitive effect of prospective gains. Whether the appetitive dimension of gambles eventually dominates automatic fear responses may then depend on the potentiation of emotional responses and on the efficiency of downstream cognitive control, which would explain why the SMG, dmPFC, dlPFC and vlPFC cortices mediate the effect of gain on decisions. This is also in line with our analysis of inter-individual differences of loss aversion, which shows that peoples’ loss aversion increases when the indirect effect size (of gains on accept decisions) in the left dlPFC pathway increases. This is because a strong involvement of the dlPFC pathway may signal inefficient cognitive control (Braver et al., 2010; Poldrack, 2015), which would result in loss aversion worsening.

Although quite self-consistent and elegant, this interpretation really relies on the “native” causal interpretation of brain-behavior mediation analysis. So what if we had not found support for this causal scenario with our I/O test statistics? In fact, the existing literature may also be queried to find past evidence that may be more compatible with the alternative causal interpretation of brain-behavior mediation analysis. For example, beyond its well-known implication in language processing, the right SMG has been shown to be involved in somatosensory perception (Ben-Shabat et al., 2015; Tunik et al., 2008). Under this perspective, evidence for *b* ≠ 0 (or, equivalently, *d* ≠ 0) may be interpreted in terms of low-level perceptual representations of (motor?) action plans. We note that the experimental design is compatible with this interpretation because the spatial arrangement of accept/reject responses is not randomized over trials (Chen, 2014). This highlights the need for developing approaches that reduce the causal ambiguity of simple brain-behavior mediation analyses.

## Appendix A: OLS estimators of path coefficients

Recall that the second line of Equation 2 can be re-written as:

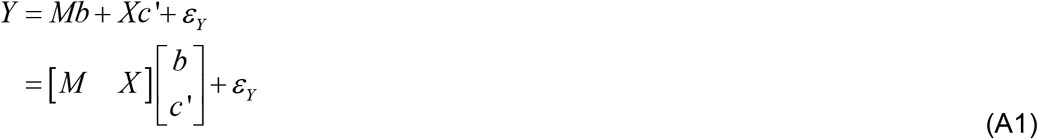

The OLS estimators of *b* and *c*′ path coefficients are thus given by:

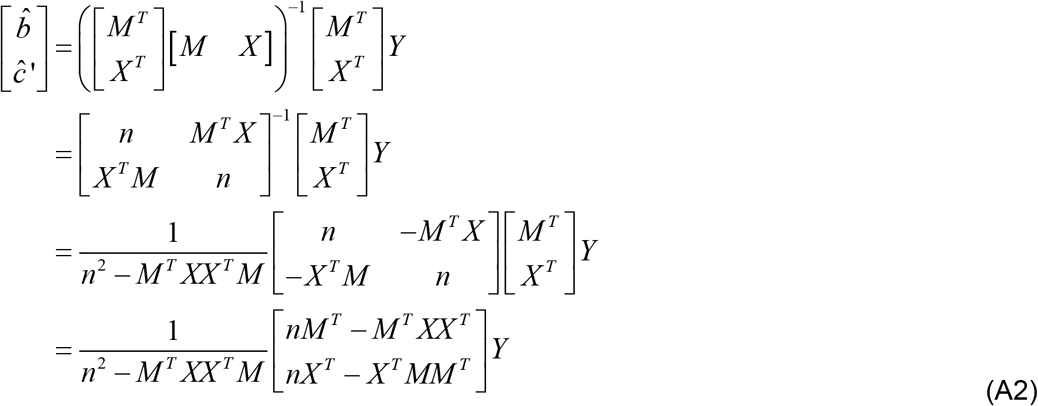

where the third line derives from the analytical formulation of 2×2 inverse matrices.

Now recall that 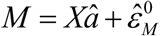 with 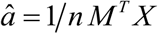. The estimator of path coefficients *b* and *c* ‘ thus writes:

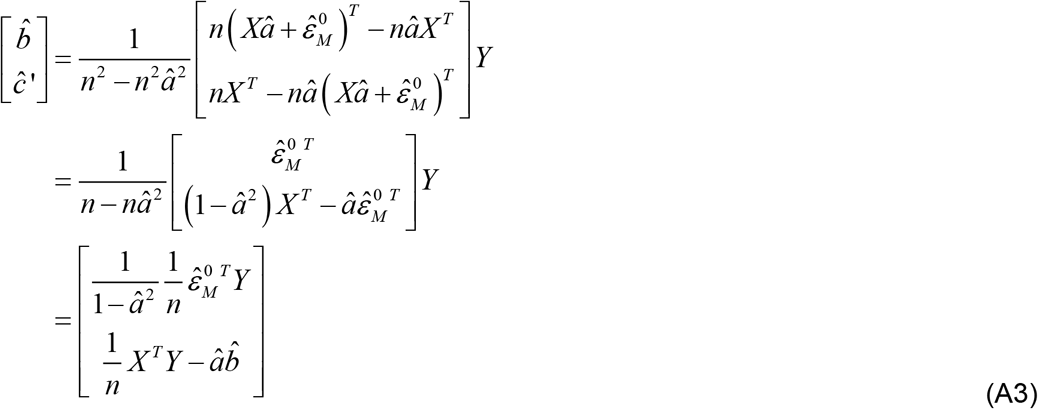

This completes the derivation of path coefficients’ estimates.

## Appendix B: Sobel’s test

In what follows, we summarize the derivation of Sobel’s mediation test.

First, recall that, given Equations 4 and 5, both 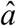 and 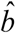 follow gaussian distributions, namely: 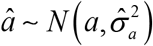 and 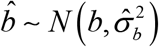. Sobel’s approach effectively reduces to a Laplace approximation of the distribution of the product 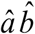 of path coefficients’ estimates.

Let 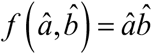 be the function that maps the pair of path coefficient estimates to their product. One can approximate 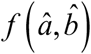 using a first-order Taylor expansion in the neighborhood of some arbitrary point (*a*_0_, *b*_0_):

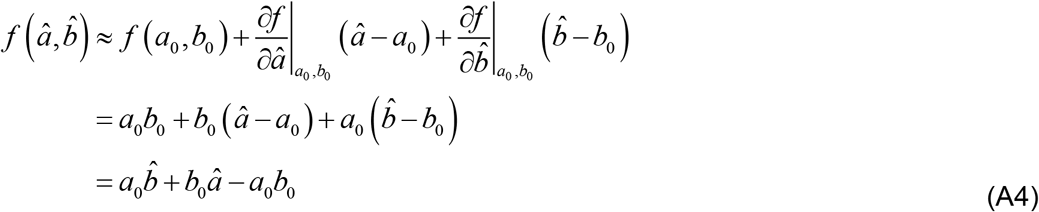

If we choose to use the above Taylor expansion in the neighborhood of the unknown true values of path coefficients (*a*_0_ ≡ *a*, *b*_0_ ≡ *b*), then Equation A4 provides us with a Laplace approximation to the first two moments of the bivariate product 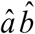:

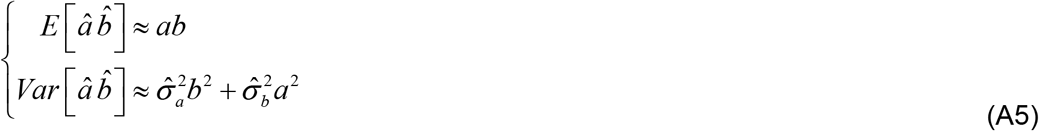

The Sobel test directly relies on this approximation to form a pseudo z-score 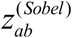 for the strength of the indirect path, as follows:

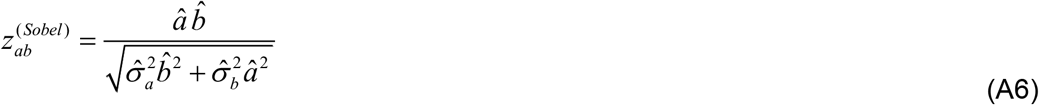

where the unknown path coefficients have been replaced by their OLS estimates. Note that 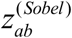 is invariant under arbitrary rescaling of *X*, *Y* and/or *M*. Under the null *H*_0_ : *ab* = 0, 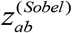 approximately follows Student’s probability density function with appropriate degrees of freedom.

We note that this approximation will be quite tight away from the diagonal lines 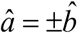, where the product 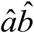 will start to behave as a quadratic function. But Sobel’s approximation error will grow quicker than estimation errors on path coefficients.

One can also show that Sobel’s test statistics is always smaller than conjunctive’s test statistics:

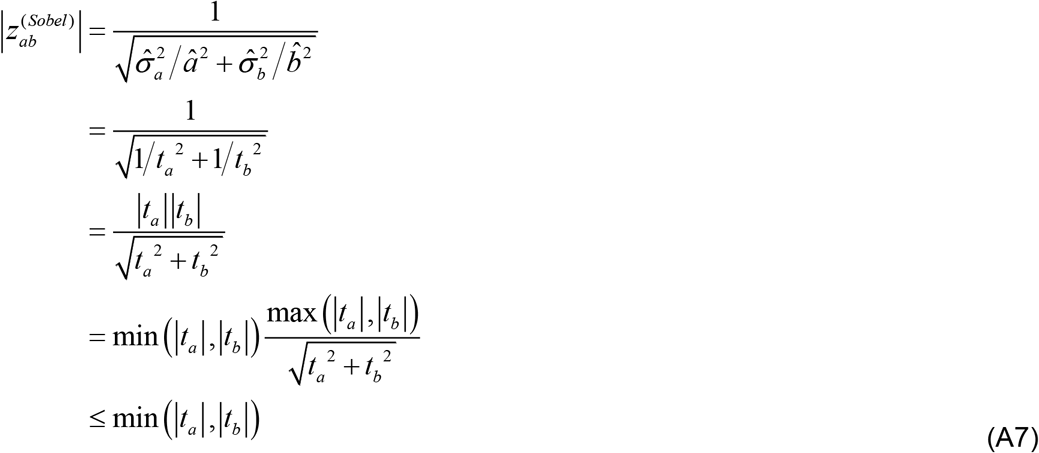

where 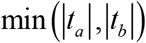 is the conjunctive test statistics (cf. Equation 9).

## Appendix C: Dealing with contrasts on experimental conditions

So far, we have only considered simple independent variables *X*. However, a typical experiment includes more than one condition or factor, and the question of interest might be best framed in terms of mediating the effect of a linear combination of independent variables. In other terms, we want to generalize classical mediation analyses of the sort implied by Equation 1 to *contrasts* of experimental factors.

Without loss of generality, let us consider an experimental design with 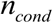 conditions, which are encoded through a *n × n_cond_* design matrix **X**. Typically, the entries of **X**’s columns would be zero everywhere, except at trials that belong to the corresponding condition (where their value would be one). Replacing *X* with the design matrix **X** in Equation 2 induces the following two-fold lieanr regression model:

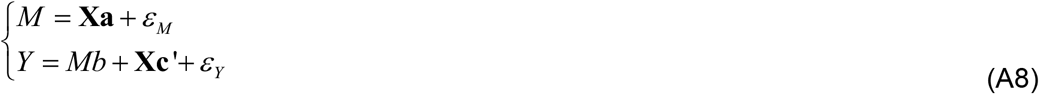

where **a** and **c**’ are now *n_cond_* ×1 vectors of regression coefficients that encode the condition means. In this context, most experimental questions of interest are framed in terms of contrasts on path coefficients **a**. So how can one ask whether *M* mediates the effect of an arbitrary contrast on path coefficients?

Two cases may arise. In the simplest case, one would deal with single contrasts. Let **w** be an arbitrary *n_cond_* vector of contrast weights. For example, in a typical 2×2 factorial design, **w** = [1 −1 −1 1] would be capturing the interaction between the two factors. Single contrasts of this sort do not require any specific adaptation of mediation analyses, because (**w**^*T*^**a**) is a scalar, and its OLS estimate has a known fixed-form distribution under the null. In turn, asking whether single contrasts are mediated by *M* reduce to testing whether, (**w**^*T*^**a**)*b* ≠ 0, which can be done using either the indirect or conjunctive approaches described above. Slightly more subtle is the case of multiple contrasts, as induced by global null hypotheses tests. For example, let us consider an experimental design with three conditions. In analogy to ANOVA, we wish to test for the mediation of *any* difference between the conditions. The corresponding contrast of interest **w** is now a 3×2 matrix of weights, and **w**^*T*^**a** becomes a 2×1 vector. Outside the context of brain-behavior mediation analysis, assessing the statistical significance of such a contrast would be performed using an F-test, for which p-values can be derived analytically (Friston et al., 1995). When using the conjunctive approach, this poses no problem, as one would simply compute the p-value of the resulting minimum F-statistics. The indirect approach is more difficult to adapt here. In principle, one would first have to partition the design matrix 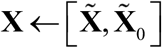 into subspaces respectively spanning the contrast of interest 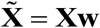 and the effects of no interest 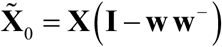. Then, one would remove the effects of no interest from both the mediator and the dependant variable. Finally, one would have to test whether *any* indirect path induced by the ensuing columns of 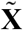 is significant. The latter issue is not entirely trivial, but can be solved using the *minimum p-value* approach (Friston et al., 1999; Nichols et al., 2005) of conjunction analysis.

## Appendix D: group-level random-effect analysis

Let us now consider the specific issue of experiments performed with multiple subjects. For example, let us assume that each subject participates in an experiment consisting of multiple trials, such that Equation 2 describes the relationship existing between *X*, *M* and *Y* across trials, at the subject-level. We now want to ask whether there is a mediated effect that is consistent across subjects, at the group-level. This calls for mixed-effects analyses, which essentially assume that subject-level path coefficients are sampled from a parent (population) distribution whose mean we wish to infer on. This can be efficiently performed using a summary statistics approach (Friston et al., 2005b; Holmes et al., 1998), whereby one first estimates subject-level effects (here, 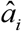 and 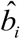, where i∈[1,…*n*] is the participant’s index), and then report these for a random-effect analysis at the group-level.

Similarly to subject-level analysis, both conjunctive and indirect approaches are possible here. Let *μ_a_* and *μ_b_* be the unknown population mean of *a* and *b* path coefficients, respectively. At the group-level, the null hypothesis of mediation analysis can be written as follows:

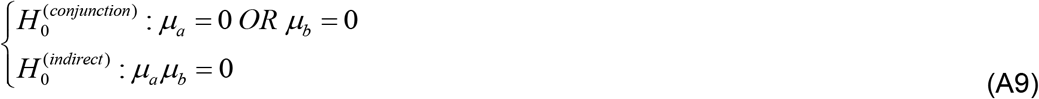

where 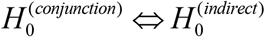 as before.

The conjunctive approach then simply reduces to testing whether both group-mean estimates 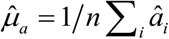 and 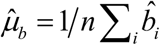 differ from zero, which can be tested using the p-value for the minimum statistic.

The indirect approach relies on testing whether the group-mean of the indirect effect differs from zero. According to the central limit theorem (Rosenblatt, 1956), the distribution of the average product 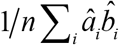 will quickly tend towards a Gaussian distribution. However, any non-zero covariance between path coefficients will bias the inference, because 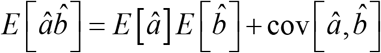 (Kenny et al., 2003). In other terms, even if the null hypothesis is true, covarying fluctuations in path estimates may significantly differ from zero. This is why the indirect approach should rather rely on testing the product 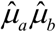 of group-mean estimates. This can be done using either parametric (cf. Sobel, Airoian or Goodman test statistics) or non-parametric (cf. M3 bootstrap) approaches.

## Appendix E: Causal impact of neural noise

The non-trivial impact of neural noise is not a feature of univariate linear brain-behavior mediation models. In fact, one can show that this generalizes to any form of brain-behavior. In what follows, we rely on an information-theoretic framework that was developed for addressing mediation claims, irrespective of the mathematical form that the mediation model may take (Pearl, 2012). The only requirement here, is that of a causal cascade from *X* to *M* and *Y*, and from *M* to *Y* (cf. directed acyclic graph in Figure 2, left panel).

Let *IE_XX_*,(*Y*) be the expected impact of the mediator variable on the behavior, under a virtual change of the manipulation (from *X* = *X* to *X* = *x′*, see Equation 9 in Pearl, 2012):

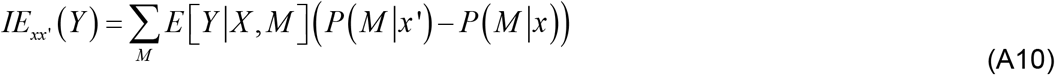

where *P*(*M|X*)is the conditional distribution of the mediator variable. Note that, in Equation A10, the causal relationships between *X*, *M* and *Y* are implicitly absorbed in conditional distributions. In brief, *IE*_xx_,(*Y*) measures the strength of the indirect effect of *X* onto *Y*, i.e. it serves as a summary statistics for significance tests of (possible multivariate and nonlinear) mediated effects.

Now, when there is no neural noise, the mediator variable brings no additional information on the behavior, i.e. *E*[*Y*|*X, M*] ≈ *E*[*Y*|*X*]. In turn, the mediator’s impact *IE*_xx′_(*Y*) becomes negligible: 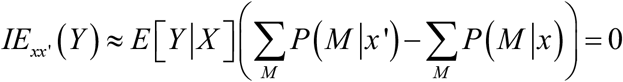. In other terms, when the mediator brings no additional information on behavior, it cannot be detected.

Conversely, when neural noise dominates, the mediator is effectively independent from the manipulation, i.e.: *P*(*M*|*X*′)≈ *P*(*M*|*X*)≈ *P*(*M*). It follows that, here again: 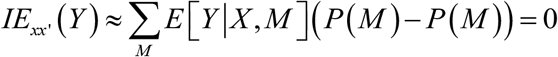. In other words, when the manipulation brings no or little information on the mediator, no mediation can be detected.

In conclusion, mediated effects can only be detected for intermediate neural noise magnitudes, irrespective of the mathematical form of the brain-behavior mediation model.

## Appendix F: Equivalence of causal interpretations of mediation analysis

In what follows we give a proof of (i) Equation 12 in the main text, and (ii) equality of t-statistics of “native” and “swapped” path coefficients.

First one can use the expressions of their OLS estimates to derive the ratio of the two path coefficients (see Appendix A):

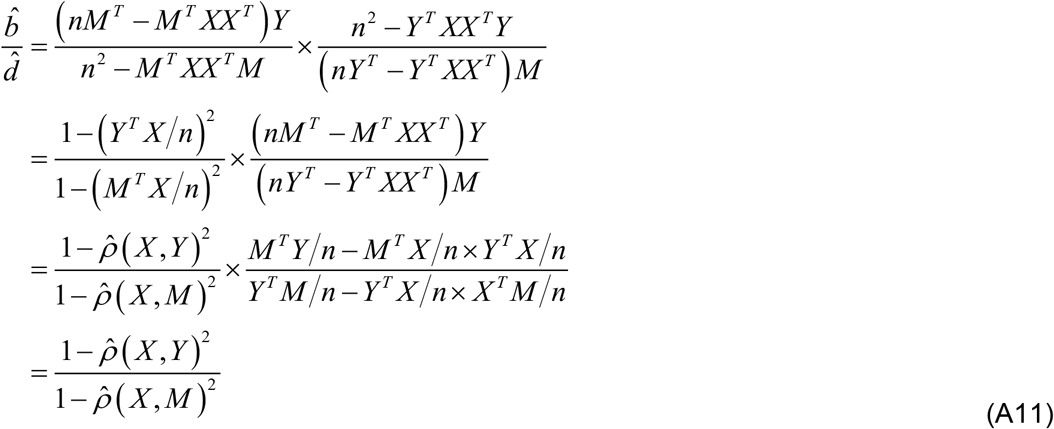

where 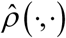 is the sample correlation between arbitrary vectors.

Now, recall that, in Equations 1–2, (i) all mediation variables are z-scored and (ii) residual estimates are, by construction, orthogonal to the variable *X*. Therefore :

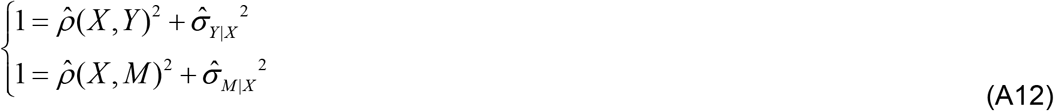

This concludes the demonstration of Equation 12 of the main text 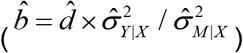.

Now let us prove the equality of t-statistics of “native” and “swapped” path coefficients.

Using the definition of these test statistics we have:

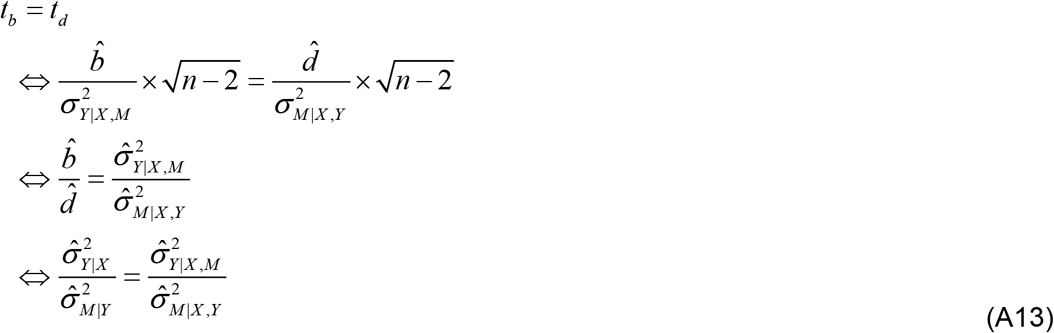

Now recall the iterative decomposition of the determinant of a gram matrix: det ([*A, v*]^*T*^[*A, v*] = det (*A^T^A*) × (*v^T^v − v^T^A*(*A^T^A*)^-1^ *A^T^v*) where *A* and *v* are arbitrary matrix and vectors, respectively (Csató and Opper, 2003). This yields:

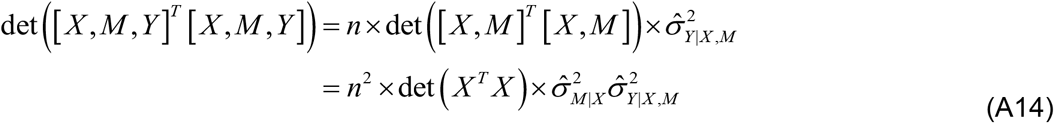

Similarly, we have:

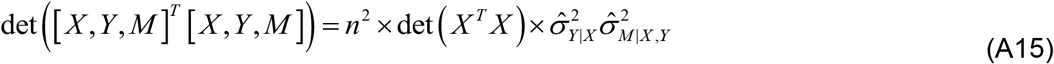

Lastly, because the order of the matrix’s columns leaves the determinant unchanged, Equations A14 and A15 are identical. This implies that 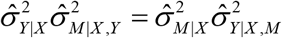, which concludes our proof.

## Notes

https://openneuro.org/datasets/ds000053/versions/00001

### Competing Interest Statement

The authors have declared no competing interest.

